# Coupling of protein condensates to ordered lipid domains determines functional membrane organization

**DOI:** 10.1101/2022.08.02.502487

**Authors:** Hong-Yin Wang, Sze Ham Chan, Simli Dey, Ivan Castello-Serrano, Jonathon A. Ditlev, Michael K. Rosen, Kandice R Levental, Ilya Levental

## Abstract

During T-cell activation, the transmembrane adaptor Linker of Activation of T-cells (LAT) forms biomolecular condensates with Grb2 and Sos1, facilitating signaling. LAT has also been associated with cholesterol-rich condensed lipid domains. However, the potential coupling between protein condensation and lipid phase separation and its role in organizing T-cell signaling were unknown. Here, we report that LAT/Grb2/Sos1 condensates reconstituted on model membranes can induce and template lipid domains, indicating strong coupling between lipid- and protein-based phase separation. Correspondingly, activation of T-cells induces protein condensates that associate with and stabilize raft-like membrane domains. Inversely, lipid domains nucleate and stabilize LAT protein condensates in both reconstituted and living systems. This coupling of lipid and protein assembly is functionally important, since uncoupling of lipid domains from cytoplasmic protein condensates abrogates T-cell activation. Thus, thermodynamic coupling between protein condensates and ordered lipid domains regulates the functional organization of living membranes.

**SUMMARY:** Membrane-associated protein condensates couple to ordered membrane domains to determine the functional organization of T-cell plasma membranes

## MAIN TEXT

Spatial compartmentalization is a ubiquitous feature of living systems, with all life on Earth compartmentalized by lipid membranes. Membranes can be further laterally sub-compartmentalized by self-organizing lipid domains. For example, the intrinsic capacity of sterols and tightly packing lipids to preferentially associate into liquid-ordered phases can produce liquid domains in biomimetic systems (*1-3*), isolated plasma membranes (PMs) (*4, 5*), and yeast vacuoles (*6, 7*). Micrometer-scale ordered membrane domains (or “rafts”) have not been directly imaged in living mammalian cells, but accumulating evidence supports the involvement of nanometer scale, dynamic lipid domains in signaling and trafficking (*8*).

A conceptually analogous organizing principle is cytoplasmic compartmentalization via biomolecular condensates, which concentrate molecules in the absence of an encapsulating membrane (*9, 10*). Similar to lipid self-organization (*11*), some condensates form through weak, multivalent interactions between biomacromolecules, which drive liquid-liquid phase separation to produce dynamic mesoscale compartments (*10*). Condensates have become extensively implicated in cellular functions, including embryonic development (*12*), synaptic organization (*13, 14*), nuclear organization, gene regulation (*15, 16*), and signaling at the plasma membrane (PM) (*17-19*). A prominent example of the latter is the PM module consisting of LAT and two cytoplasmic adaptors Grb2 (growth factor receptor-bound) and Sos1 (son of sevenless), which links T-cell immune receptor engagement with downstream pathways for activation (i.e. proliferation, cytokine secretion, etc.) (*20*). Interactions between LAT/Grb2/Sos1 produce liquid condensates in reconstituted and living systems (*18, 19*), which affect signaling by (a) concentrating reactants (*18, 21*); (b) excluding negative regulators, e.g. the phosphatase CD45 (*18*); (c) coupling to cytoskeletal dynamics (*19*); and (d) kinetically proofreading activation (*21*).

A central outstanding question concerns the biophysical and functional coupling between membrane lipid domains and cytoplasmic condensates (*22, 23*). Lying near phase coexistence boundaries, both the PM (*24, 25*) and cytoplasm (*9*) appear poised for large-scale structural rearrangements, such that phase separation of one could produce significant responses in the other. We hypothesized that LAT could produce such coupling, because it participates in both cytoplasmic condensates via its disordered cytoplasmic tail (*18, 19*) and ordered lipid domains via its transmembrane helix (*26-29*). Through a combination of *in vitro* reconstitution and cellular experiments, we show that LAT condensates are thermodynamically coupled with ordered membrane domains during T-cell activation, providing direct evidence of convergence between phase separation of membrane lipids and protein condensation. We further show that this coupling is functionally important, as uncoupling abrogates cell activation downstream of T-cell receptor engagement. These observations indicate that protein condensates can regulate the functional organization of lipid membranes, and conversely, that lipid phase separation can potentiate membrane protein condensation.

### Mutual templating between protein condensates and membrane domains in vitro

Investigations of membrane-associated protein condensates have often relied on planar lipid bilayers formed by fusion of liposomes onto solid supports (*18, 19*). This method is not amenable for studying ordered membrane domains because cholesterol-rich mixtures fuse poorly (*30*) and domain properties are severely affected by the solid substrate (*31, 32*). To overcome these limitations, we generated phase-separating lipid multi-bilayers by spin-coating mixtures of cholesterol, saturated phosphatidylcholine (DPPC), and unsaturated phosphatidylcholine (DOPC), which form dynamic, temperature-reversible liquid-ordered (Lo) and -disordered (Ld) domains (*33*) (Fig S1). The phosphorylated intracellular domain of LAT (pLAT) was bound to the topmost leaflet of the multi-bilayer through lipids with a His-chelating headgroup (e.g. DSIDA, which concentrates in Lo domains) (*34*) (Fig S2).

As previously shown (*18, 19*), pLAT coupled to a DOPC membrane phase separates to form micron-sized condensates within minutes of introduction of Grb2+Sos1 (Fig S3) (throughout the text we invoke phase separation for in vitro experiments where this mechanism of protein assembly has been clearly established and the more generic ‘condensate’ for cellular results where the mechanism of assembly is less definitive). On more biomimetic, phase-separated membranes, DSIDA-anchored pLAT was uniformly distributed in the Lo phase in the absence of Grb2+Sos1 (Fig 1A, top), evidenced by its segregation from the Ld domain marker (TR-DHPE) (see figure legends and Supplementary methods for lipid and protein concentrations). Addition of Grb2+Sos1 induced pLAT coalescence into large condensates that were exclusively overlying the Lo regions (Fig 1A, bottom). These condensate-rich regions existed alongside condensate-poor Lo regions, consistent with three-phase (condensate-rich Lo, condensate-poor Lo, and Ld) coexistence predicted by recent theory (*35*).

**Figure 1.**
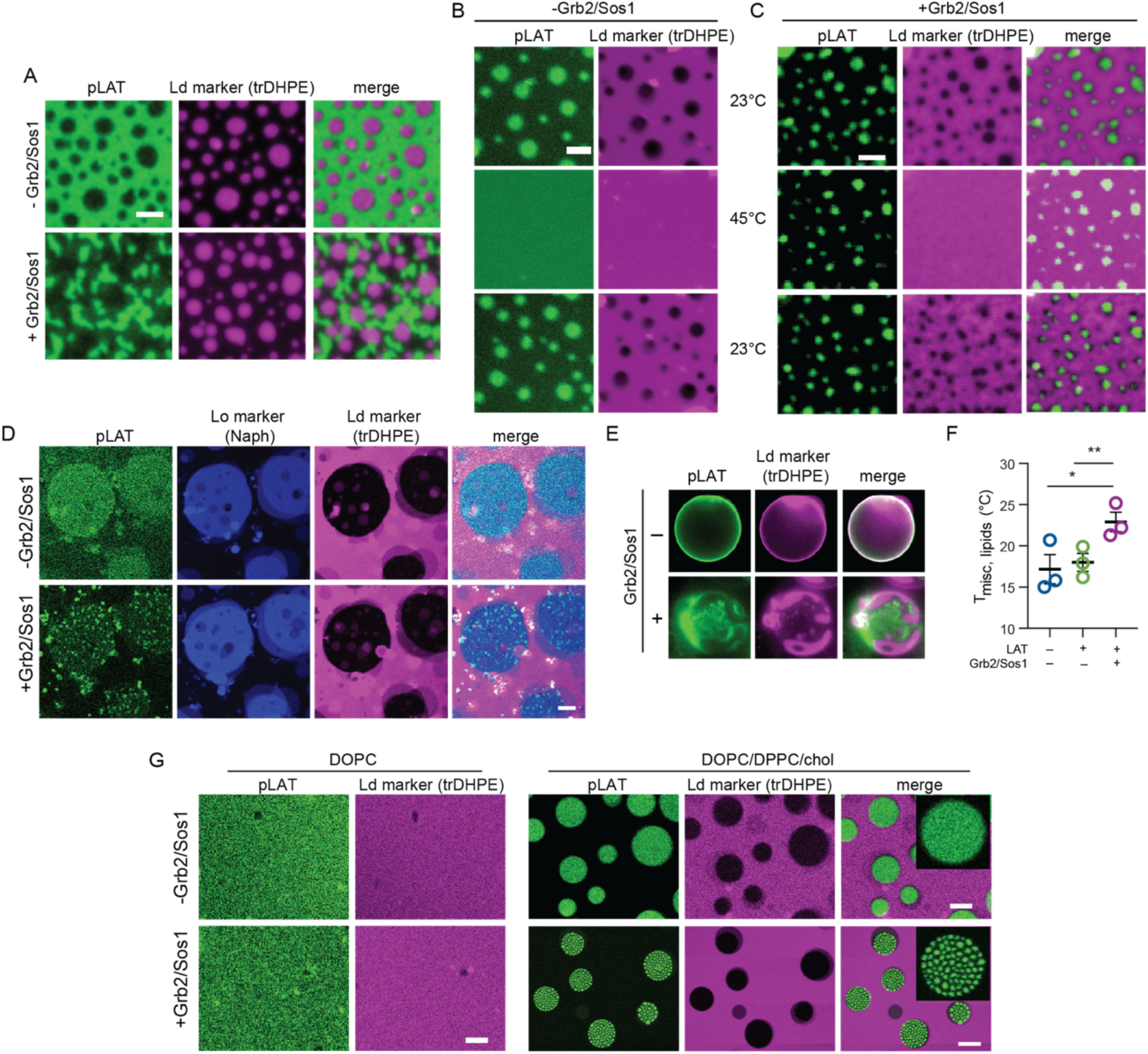
*in vitro* coupling between protein condensates and membrane domains. (A) pLAT is uniformly distributed in Lo domains in a phase separated membrane when conjugated to DSIDA. Addition of Grb2/Sos1 produces protein condensates exclusively on top of Lo domains, i.e. completely excluded from Ld domains (marked by trDHPE). (B) Lo domains enriching pLAT are present at 23°C and disperse at 45°C, then reappear randomly after cooling. (C) Condensates recruit nascent Lo domains. LAT/Grb2/Sos1 condensates form in small Lo domains in majority-Ld membranes. Lipid domains are dissolved by increasing temperature above the miscibility transition threshold (∼37°C for this composition, see Supp Methods for detailed lipid compositions); pLAT condensates were not notably affected at these temperatures. Cooling membranes below the transition temperature induces Lo domain formation exclusively beneath protein condensates. (D) pLAT was bound to both phases by 2% DP-NTA and 1% DO-NTA in the bilayer, yielding ∼2-fold Lo enrichment of LAT. Adding Grb2/Sos1 produces condensates exclusively in the Lo phase (labeled by naphthopyrene, blue). (E) Protein condensates can induce lipid phase separation in GUVs. (F) Protein condensates enhance lipid phase separation, evidenced by significantly increased T_misc_ (temperature at which 50% of GUVs show lipid phase separation). (G) Lipid phase separation facilitates protein condensation. Low concentrations of pLAT (20 nM), Grb2 (100 nM), and Sos1 (100 nM) do not produce pLAT condensates on DOPC membrane (left panel), whereas the same protein mixture undergoes Lo-confined condensation on phase separated membranes (right panel). Scale bars are A/C/D/E/G = 5 µm, B = 2 µm.

At this membrane composition, Lo domains comprise most of the bilayer area; to determine whether condensates could recruit Lo domains, the abundance of unsaturated DOPC was increased to produce Ld-majority bilayers (Fig 1B). In these, pLAT was confined to small Lo domains (Fig 1B, top) and addition of Grb2+Sos1 did not affect its superficial appearance (Fig 1C, top). However, Grb2+Sos1 did induce LAT condensation, as evidenced by the persistence of the pLAT clusters upon melting of the underlying membrane domains at 45°C (Fig 1C, middle). This behavior contrasts with that of pLAT alone, which disperses together with domains at 45°C (Fig 1B, middle). When membranes were again cooled below the miscibility transition temperature (T_misc_), Lo domains reappeared exclusively beneath the pLAT/Grb2/Sos1 condensates, but randomly on bilayers containing only pLAT (Fig 1B-C, bottom). These observations were mirrored when pLAT was instead attached to the Ld phase (Fig S4). Thus, lipid domains and LAT/Grb2/Sos1 condensates can mutually template each other’s localization and morphology.

We next tested how condensation would affect pLAT organization when monomers were not confined to a single phase, but rather partitioned more like in isolated plasma membranes, where it is enriched by 20-50% in the raft phase (*26, 27, 36*). To that end, pLAT was coupled to bilayers via a mixture of saturated (DP-NTA) and unsaturated (DO-NTA) tail lipids. At a ratio of 2:1 DP-NTA:DO-NTA, pLAT partitioned to both phases with a moderate preference for the Lo phase (K_p,Lo_ = 2.1±0.4), Fig 1D, S5A-B), as in natural systems. Strikingly, LAT/Grb2/Sos1 condensates were observable exclusively in the Lo phase (Fig 1D). Even when condensates formed in regions where Ld phases dominated, they appeared to induce the formation of small Lo domains (Fig S5A). We hypothesize that this high affinity of condensates for Lo domains is driven by oligomerization: the partitioning free energy of individual LAT monomers (into Lo) is additive, meaning that partition coefficients multiply such that K_p,oligomer_ goes as K_p,monomer_^N^, where N=oligomer number. Thus, even for weakly partitioning monomers, this exponential dependence on oligomer number would dramatically enhance partitioning for large oligomers like condensates.

### Reciprocal stabilization between protein condensates and membrane domains

To investigate potential thermodynamic coupling between protein phase separation and lipid phase separation, we measured the stability of membrane domains in Giant Unilamellar Vesicles (GUVs). GUVs composed of DOPC, DPPC, and 40% chol were not phase separated (homogeneously distributed fluorescent lipids and DSIDA-coupled pLAT) at 23°C (Fig 1E). As on supported bilayers, addition of Grb2+Sos1 coalesced pLAT into condensates, but also induced phase separation in the lipids, with complete segregation between an Ld phase marker (trDHPE) and protein condensates (Fig 1E, bottom). These observations suggest that protein condensates can stabilize membrane domains, consistent with reports that clustering of membrane components can induce domain formation in cholesterol-containing membranes (*25, 37, 38*). To define the magnitude by which protein condensates potentiated membrane phase separation, we measured condensate-dependent phase separation in GUVs (Fig 1F and S6). At low temperatures, GUVs separate into coexisting Lo/Ld domains, which ‘melt’ (i.e. become miscible) at higher temperatures. The temperature at which 50% of vesicles are phase separated is defined as the miscibility transition temperature (T_misc_), which in the absence of LAT was 17°C. pLAT conjugated to the Lo phase via DSIDA did not affect the thermotropic phase transition, whereas condensates induced by Grb2+Sos1 increased T_misc_ to 22°C (Fig 1F and S6), indicating robust regulation of lipid phase transition by LAT condensates, as also recently reported (*39*). Similar effects were observed when pLAT was initially coupled to both phases via the mixture of saturated/unsaturated-lipid-NTA, as in Fig 1D (Fig S5B-C). Thus, protein condensates enhance membrane phase separation, likely via clustering domain-associated components, as previously observed for membrane-bound actin filaments (*40-43*), glycolipids (*25*), and other condensing proteins (*39, 44*).

Inversely, we predicted that enrichment of LAT into ordered domains, as believed to occur in living cells (*27, 29*), would increase its propensity to form condensates. We tested this prediction by lowering the concentration of LAT/Grb2/Sos1 to a regime where no condensates are observable on a uniform Ld membrane (Fig 1G, left). The same protein mix applied to a phase-separated membrane robustly induced condensates confined within Lo domains (Fig 1G, right, Fig S7). Thus, protein condensates and membrane domains can reciprocally stabilize each other, i.e. they are thermodynamically coupled.

### Grb2 condensates recruit ordered membrane environments in cells

We next investigated coupling of LAT/Grb2/Sos1 condensates with membrane domains in living cells. To induce LAT condensates, Jurkat T-cells were activated by seeding onto coverslips coated with OKT3, an α-CD3 antibody. Crosslinking of CD3 triggers a signaling cascade that converges on LAT phosphorylation (*45*), which in turn recruits Grb2 and Sos1 to form cytoplasmic condensates analogous to those observed *in vitro* (*18*) (Fig 2A-B). To evaluate the coupling of these condensates with membrane domains, we employed genetically encoded probes that differentially partition to membrane phases (*26, 27, 36*). As reporters of ordered membrane regions, we used the transmembrane domain (TMD) *α*-helix of LAT (raft-TMD) (*26, 27, 36*) or a saturated lipid-anchor (GPI, glycophosphatidylinositol). To mark disordered regions, we used a TMD consisting of 22 Leu residues (nonraft-TMD) (*26, 27*). All probes were fused to fluorescent proteins for visualization and validated for expected raft affinity (Table S1). Grb2 condensates were coincident with regions of clear enrichment of the two raft probes (raft-TMD and GPI) and were depleted of the nonraft-TMD (Fig 2C-F). The magnitudes of enrichment/depletion were consistent with previous reports for enrichment of raft-associated markers around activated immune receptors by super-resolution localization microscopy (*46*). Similar sorting of raft markers with Grb2 condensates was observed by inducing condensates on non-activating surfaces (ICAM1) with the phosphatase inhibitor pervanadate (Fig S8). Lipid fluorophores that selectively enrich in Lo or Ld domains were also sorted in accordance with raft enrichment at condensates. (Fig S9). Thus, activation-induced protein condensates are co-localized with raft-like membrane environments in living Jurkat T-cells, in agreement with reconstitution experiments.

**Figure 2.**
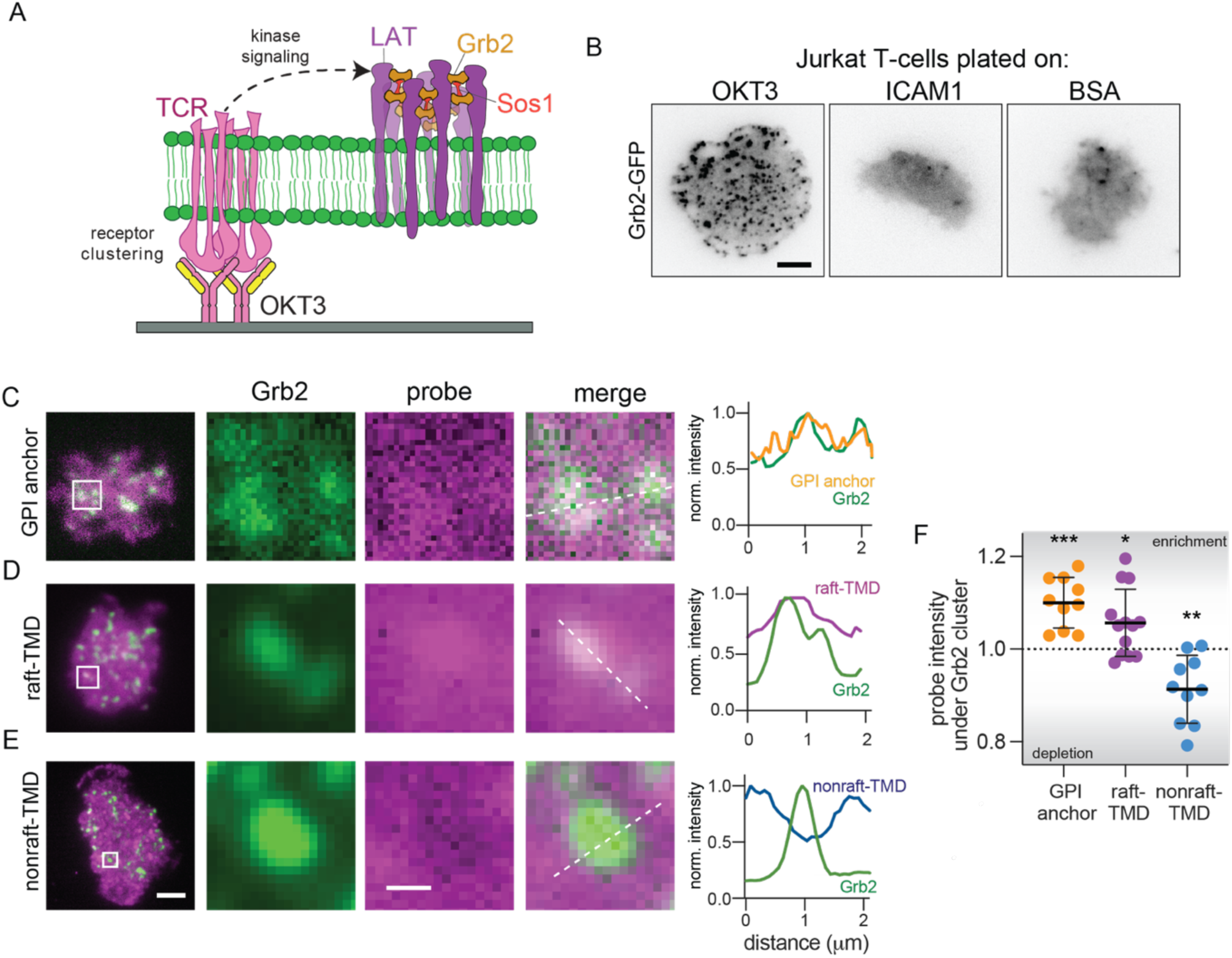
*in situ* recruitment of raft marker proteins to Grb2 condensates in cells. (A) Schematic of LAT condensate formation in activated Jurkats: T-cell receptor (TCR) clustering by OKT3 induces signaling that leads to phosphorylation of LAT and multivalent recruitment of Grb2/Sos1 to produce liquid condensates. (B) TIRF imaging of Grb2-rich condensates formed in OKT3-activated Jurkats, but not in non-activated cells (BSA- or ICAM1-coated coverslip) Scale bar is 5 µm. (C-E) TIRF images of Grb2 condensates relative to GPI-GFP, raft-TMD, and nonraft-TMD. Scale bar is 5 µm. Enlarged images of white squares demonstrate recruitment of raft-TMD / GPI-GFP and exclusion of nonraft-TMD under Grb2 condensates. Scale bar is 0.5 µm. (right) line scans showing probe enrichment under Grb2 condensates. (F) Quantification of relative enrichment in cells imaged at room temperature (imaging at 37°C gave similar results, Fig S10). Each point represents the mean enrichment (>1) or depletion (<1) of probes under Grb2 condensates relative to adjacent region for individual cells across 3 independent experiments. Each cell included >10 Grb2 condensates. ***p<0.001, **p<0.01, *p<0.05 for difference from 1 (no enrichment/depletion) of means of individual cells.

### Cooperative recruitment between Grb2 condensates and micron-sized membrane domains in live cells

Raft probe enrichments were relatively subtle and only observable by using Grb2 condensates as fiducial markers, putatively because the probes used all have relatively low selectivity for raft domains (*26, 36*). To more clearly detect coupling, we relied on a previously described strategy to enhance probe raft affinity by oligomerization (*1, 36*). To this end, we used antibodies to crosslink an endogenous T-cell raft component, the GPI-anchored protein (GPI-AP) Thy1 (*47*). The efficacy of this approach was confirmed in cell-derived Giant Plasma Membrane Vesicles (GPMVs), where primary antibody-crosslinked Thy1 had significantly higher raft phase affinity than monomeric GPI-AP (Fig 3A-B). Further clustering of Thy1 by secondary antibodies produced large clusters that precluded accurate raft affinity measurements; however, essentially all clusters were found in the GPMV raft phase (Fig 3A).

**Figure 3.**
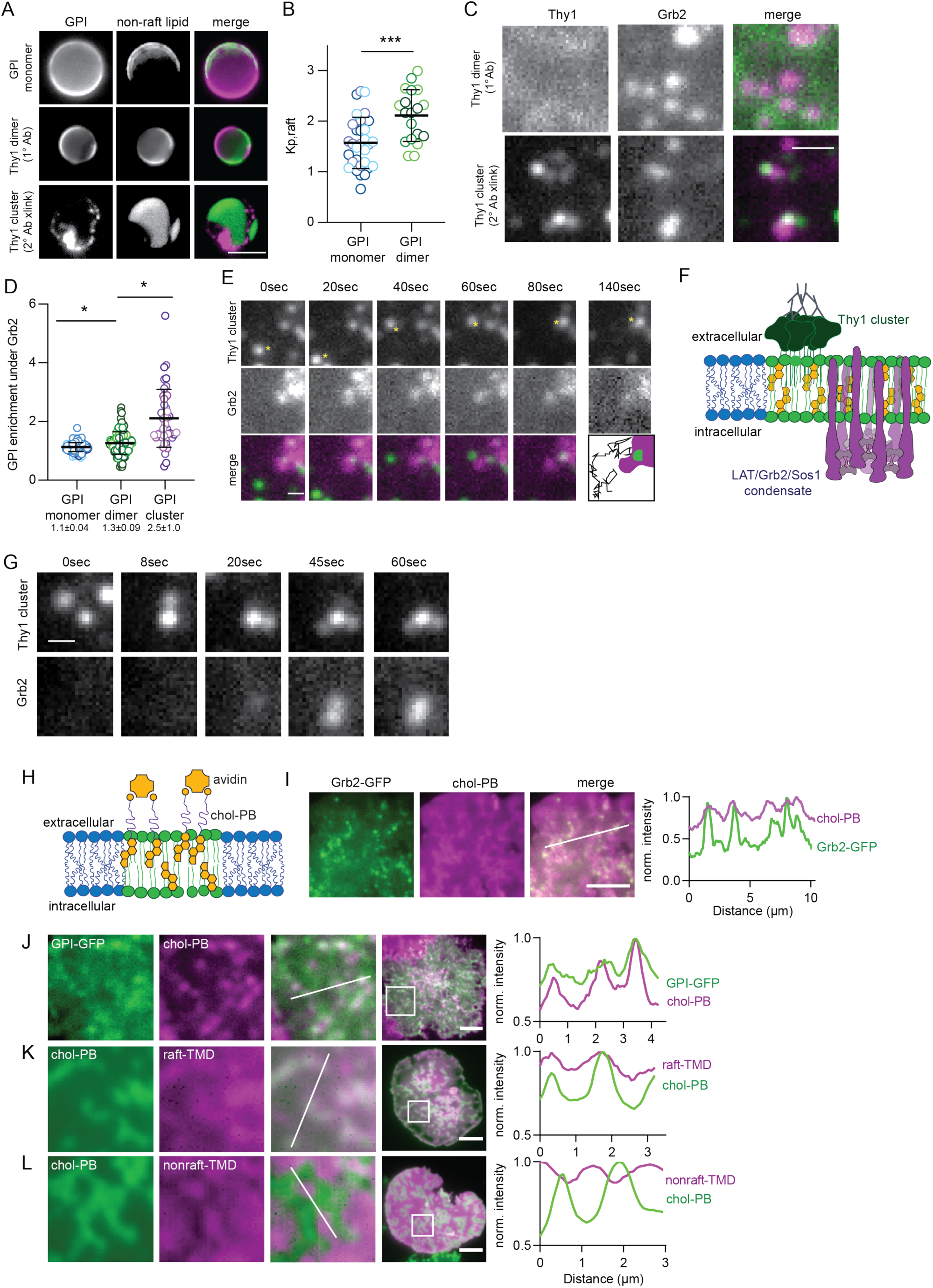
Mutual templating between Grb2 condensates and raft-like membrane domains. (A) Monomeric GPI AP (GPI-GFP) is subtly enriched in raft phase of phase separated GPMVs. Dimerization of the endogenous GPI-AP Thy1 increases raft preference. Further oligomerization of Thy1 via secondary antibodies leads to exclusively raft-associated clusters. Scale bar is 5 µm. (B) Quantification of the partitioning coefficient (K_p,raft_) showed that antibody dimerized Thy1 has a higher raft affinity than non-crosslinked GPI-AP. (C) Recruitment of oligomerized Thy1 to Grb2 condensates. Top: overlapping of Grb2 condensates with primary antibody dimerized Thy1. Bottom: overlapping of Grb2 condensates with Thy1 clusters induced by secondary antibody crosslinking. Scale bar is 1 µm. (D) Quantification of GPI-AP enrichment under Grb2 condensates enhanced by antibody crosslinking. Data points represent individual cells from at least three independent experiments; *p<0.05. (E) Time-lapse of live cell imaging showing capture/immobilization of a Thy1 cluster by Grb2 condensate. Scale bar is 0.5 µm. (F) Schematic of coupling between clustered Thy1 and LAT/Grb2/Sos1 condensate mediated by ordered membrane domain. (G) Time series showing formation of Grb2 condensate above an immobilized Thy1 cluster. Scale bar is 1 µm. (H) Schematic of cholesterol-PB oligomerization and labeling. Cells are labeled with chol-PB, then fluorescent avidin, before plating on OKT3-coated coverslips. (I) TIRF images of Grb2 condensates overlying micron-sized cholesterol-rich domains. Plot shows normalized fluorescence intensity along the line trace shown in white. (J-L) TIRF imaging reveals the colocalization of cholesterol-rich domains with raft markers GPI-GFP and raft-TMD, and exclusion of nonraft-TMD from chol-rich domains. Scale bars are 5 µm. Plots show normalized fluorescence intensity along the line traces.

Consistent with our prediction that enhancing raft affinity would amplify colocalization with condensates, dimerization of Thy1 by primary antibody significantly increased enrichment under Grb2 condensates generated by TCR activation compared to monomeric GPI-AP (Fig 3C-D). Further crosslinking by secondary antibody produced Thy1 clusters that were strongly enriched in areas juxtaposed to Grb2 condensates (Fig 3C-D). In live cells, these Thy1 clusters rapidly diffused on the extracellular surface of the PM (especially early in the activation time course) and were often observed stopping underneath Grb2 condensates (Fig 3E, Supplementary Movie 1). We conclude that the multiplicity of saturated acyl chains (previously estimated at ∼20 proteins/cluster (*48*)) in these GPI-AP clusters enhances their affinity for transbilayer ordered membrane domains (*1*), which tend to be associated with, and immobilized by, Grb2 condensates (Fig 3F). The recruitment of ordered domains by LAT/Grb2/Sos1 condensates is directly analogous to our observations in reconstituted membranes (Fig 1C). Correspondingly, after TCR activation we also observed that membrane regions marked by Thy1 clusters could nucleate Grb2 condensate formation (Fig 3G, Supplementary Movie 2). We hypothesize that these nascent condensates were induced by enrichment of LAT in membrane domains (*26, 27*) (as in the reconstitution experiment in Fig 1G), suggesting that cytoplasmic conditions are poised such that protein condensation can be induced by localized protein concentration via lipid domains.

Following a similar strategy of enhancing raft affinity via oligomerization, we evaluated the effects of crosslinking the prototypical raft lipid, cholesterol (*8*). Cholesterol was modified with a biotinylated PEG-spacer (Fig 3H), relying on a strategy validated in design of other raft probes (*49, 50*). Labeling and crosslinking with fluorescent avidin (Av647/Av488) confirmed that oligomerized cholesterol-PEG-biotin (chol-PB) enriches in the ordered phase of GPMVs more strongly than monomeric chol-PEG-FITC (Table S1 and Fig S11). In PMs of activated Jurkat T-cells, avidin-labeled chol-PB enriched in micron-sized domains (Fig 3I-L). Strikingly, Grb2 and LAT condensates were found exclusively overlying these cholesterol-rich regions (Fig 3I, S12), as small foci distributed throughout relatively larger membrane domains.

Direct microscopic observations of cholesterol-rich domains in live cells were surprising, as raft domains have largely evaded unambiguous microscopic detection (*8*). These domains were initially observed by TIRF but were also visible by epifluorescence (Fig S13) and confocal (Fig S14) imaging and were not membrane accumulations or large invaginations/deformations (Fig S13-15). Most importantly, chol-PB domains selectively recruited raft markers, including GPI-GFP (Fig3J), glycolipid-binding cholera toxin B (CTxB) (Fig S16A), and Thy1 clusters (Fig S17), all of which enrich in rafts due to saturated acyl chains (Table S1). Chol-PB domains also enriched raft-TMD and excluded nonraft-TMD (Fig 3K-L). Another non-raft TMD, from the immune cell phosphatase CD45 (Table S1), was also robustly excluded from chol-PB domains (Fig S16B). Broadly similar selective domains were also observed in Jurkat cells plated on fluid synthetic supported bilayers (Fig S18). Thus, mesoscopic raft domains are strongly coupled to Grb2/LAT condensates, revealing cooperative templating between protein condensates and membrane domains in cells, directly analogous to reconstituted systems (Fig 1A-D).

### Condensates stabilize cell membrane domains

Reconstituted condensates stabilize membrane domains *in vitro* (Fig 1E-F, S6). We hypothesized that the microscopic raft domains revealed by chol-PB were potentiated by condensation of raft-associated proteins (i.e. LAT) induced by T-cell activation. Consistently, micron-scale cholesterol-rich patterns were only observable in activated Jurkat T-cells (i.e. in presence of condensates), whereas cells plated on a non-activating (ICAM1-coated) surface were more laterally homogeneous (Fig S19A). The distribution of pixel intensities confirmed that OKT3 patterns were bimodal whereas ICAM1 were normally distributed (Fig S19B); quantification by coefficient of variation (CoV) of chol-PB intensity revealed significant differences between cell populations (Fig S19B-inset), and there were robust spatial autocorrelations in chol-PB intensity in activated Jurkat T-cells, compared to smaller scale and amplitude autocorrelations on ICAM1 (Fig S20). Thus, membrane-associated condensates reorganize the PM in live cells, enhancing the propensity to form large cholesterol-rich lipid domains (Fig 4H).

**Fig 4.**
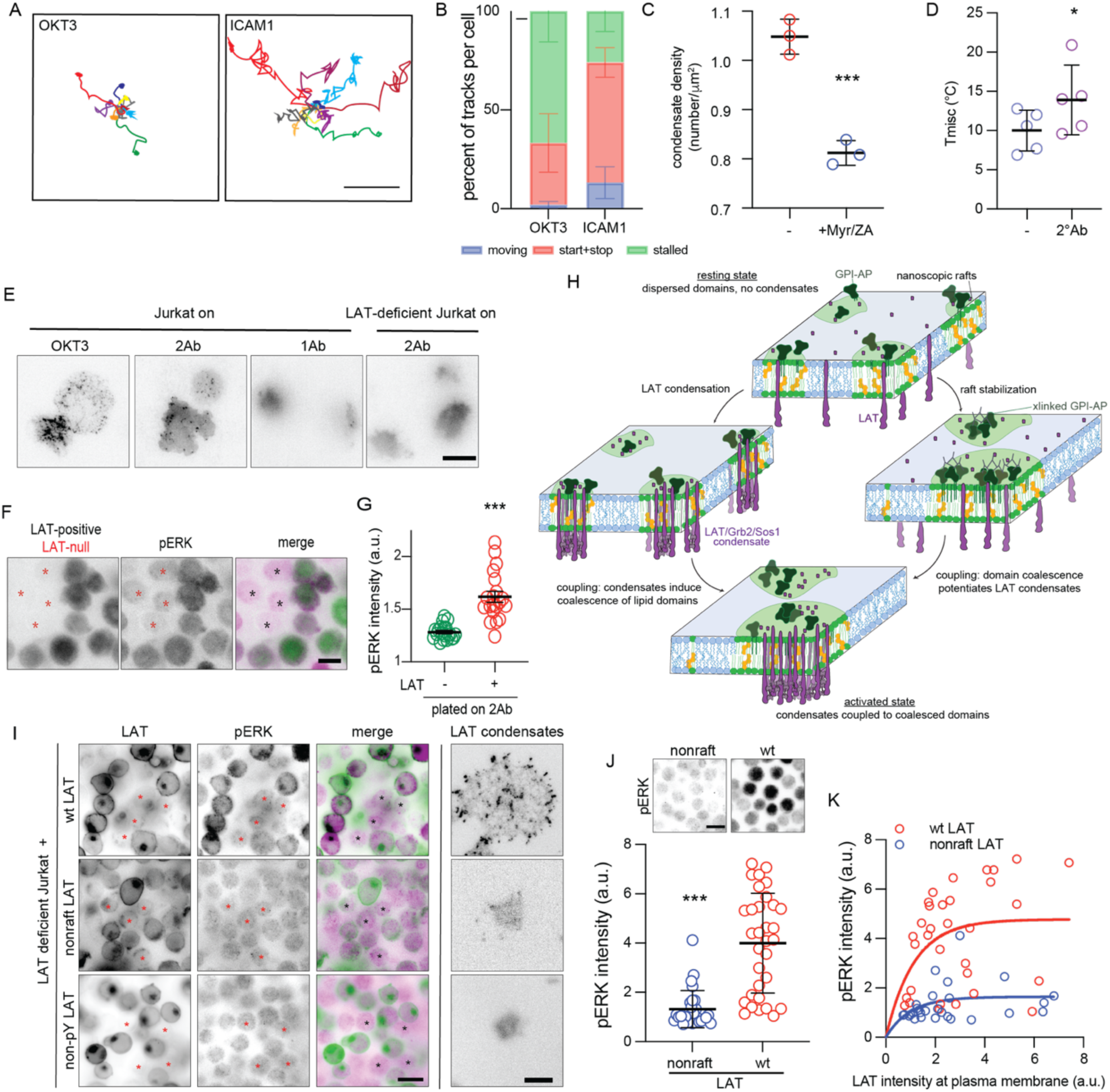
Protein condensates potentiate membrane domains. (A) 8 representative >10 sec tracks of Thy1 clusters from a cell plated on either OKT3 (left) or ICAM1 (right). (B) Trajectories of 2° antibody-crosslinked Thy1 clusters were calculated through single particle tracking (30-100 tracks/cell; five cells/condition). Each track was classified as either mobile (<20% of the track time stalled), start-stop (21-79% stalled), or stalled (≥80% stalled). Shown are percentages (mean +/-SD) of each class of track across individual cells. (C) Inhibition of raft-forming lipids perturbs Grb2 condensate formation. Grb2-mScarlet-expressing Jurkat T-cells were incubated with 25 *μ*M myriocin and 5 *μ*m Zaragozic acid for 3 days to deplete cells of lipids necessary for raft formation (e.g. sphingomyelin and cholesterol, respectively) as previously described (*54*). Shown is condensate density for 3 independent experiments with >5 cells/experiment. ***p<0.001 for t-test across experiments. (D) T_misc_ is higher in GPMVs with 2°-antibody-crosslinked Thy1, indicating increased raft stability. (E) Grb2-scarlet transfected cells imaged via TIRF 20 min after plating. 2°Ab-coated coverslips crosslink 1°Ab-labeled endogenous Thy1, which is sufficient to induce condensates in Jurkat T-cells, but not in LAT-deficient cells (JCam2.5). Scale bar is 10 µm. (F) LAT-deficient cells (marked with asterisks) were mixed with LAT-positive Jurkat T-cells (labeled with Grb2-mScarlet) and cell activation induced by Thy1 crosslinking by 2°Ab-coated coverslips was examined by pERK immunostaining. Thy1 crosslinking induced pERK, but not in LAT-deficient cells. Scale bar is 10 µm. LAT-deficient cells served as internal negative control for IF staining and imaging. (G) Quantification of pERK activation by Thy1 clustering in LAT-deficient JCam2.5 cells (green) versus LAT-expressing Jurkat cells (red). (H) Schematic model of condensate-domain coupling. (left) Clustering LAT by intracellular condensates enhances LAT recruitment into membrane domains and promotes their coalescence. (right) Clustering raft components stabilizes membrane domains to potentiate LAT condensation. (bottom) Both processes result in an activated state where membrane-associated condensates template and stabilize raft-like membrane domains. (I) Coupling of protein condensates and lipid domains through LAT is necessary for pERK activation. LAT-deficient JCam2.5 cells were mixed with cells stably repleted with either wt, non-raft, or non-pY LAT and plated on OKT3 for 10 mins. Only wt-LAT repleted cells formed LAT condensates and had pERK staining above LAT-deficient negative controls (selected LAT-deficient cells marked with asterisks). Scale bar is 15 µm (left) and 5 µm (right). (J) Quantification of pERK in raft versus non-raft LAT repleted cells. Scale bar is 15 µm. (J) pERK intensity as a function on LAT on the PM. Mean ± SD of individual cell quantifications shown in G, J-K for one representative experiment. Three independent experiments were performed with similar results.

We further explored this effect by analyzing the dynamics of antibody-crosslinked Thy1 clusters as reporters of membrane organization, with immobile clusters reflecting the presence of underlying ordered membrane domains (Fig 3E-G, as previously described (*48, 51*). Thy1 clusters were notably less dynamic in the presence of LAT/Grb2 condensates (i.e. in Jurkat T-cells activated by OKT3), stalling frequently and for long periods, compared to more mobile clusters in the absence of condensates on ICAM1-coated coverslips (Fig 4A, S21). The majority of Thy1 clusters in activated Jurkats were stalled for >80% of any individual track (minimum track length = 10 sec), whereas most clusters on ICAM1-plated cells were either mobile throughout the tracking or exhibited short, intermittent stalls (Fig 4B, S21, Supplementary Movies 3&4). These observations suggest that induction of condensates stabilized ordered membrane domains analogous to previous demonstrations of crosslinking-induced domains in model membranes (*25, 37, 46*) or actin-asters associated with GPI-rich domains in cells (*43*). This induction and recruitment of membrane domains by LAT/Grb2/Sos1 condensates may explain previous reports of selective enrichment of membrane raft markers around activated immune receptors (*46, 52, 53*).

### Cell membrane domains potentiate condensates

Since lipid domains and protein condensates are coupled in purified systems and living cells, we next tested whether perturbing membrane domains would affect LAT/Grb2/Sos1 condensates in activated T-cells. Membrane domains can be disrupted by inhibiting synthesis of raft-forming lipids using a combination of myriocin to inhibit sphingolipid synthesis and Zaragozic acid (ZA) to inhibit cholesterol synthesis. This treatment disrupts raft-like nanodomains in cells (*54, 55*) and inhibits membrane phase separation in isolated GPMVs (*56*) without off-target effects on cellular phospholipid composition (*54*). Treatment with myriocin+ZA significantly reduced condensate density in Jurkat T-cells activated by OKT3 (Fig 4C).

Conversely, crosslinking of raft components has been shown to promote and stabilize membrane domains (*25, 48, 57*) (Fig 1E-F). Consistently, crosslinking Thy1 with antibodies stabilized membrane domains in GPMV experiments, indicated by increased lipid phase separation temperature (T_misc_) (Fig 4D, Fig 4H right). Remarkably, stabilizing cell membrane domains without any other activating stimulus was sufficient to produce cytoplasmic protein condensates. Plating Jurkat cells labeled with anti-Thy1 antibodies onto 2°Ab coated coverslips to crosslink the GPI-anchored protein induced Grb2 condensate formation (Fig 4E). No condensates were observed in absence of GPI crosslinking (on 1°Ab-coated coverslips) nor in the absence of LAT (i.e. LAT-deficient Jurkat line, JCam2.5). Condensates induced by GPI-AP crosslinking were able to activate MAPK signaling, revealed by immunostaining against phosphorylated ERK (pERK) (Fig 4F-G). These results suggest that stabilizing membrane domains can induce condensate formation and downstream T-cell activation, even in the absence TCR, mediated by LAT coupling between membrane domains and condensates. Several groups have previously reported that crosslinking GPI-APs or raft glycolipids can activate T-cells without TCR ligation (*39, 48, 58, 59*). Here, we show that LAT (as a transmembrane link) is necessary and suggest a mechanism for these puzzling findings, i.e. that stabilizing rafts potentiates protein condensates (Fig 4H) that facilitate T-cell signaling (*21*).

### Uncoupling of membrane domains from protein condensates abrogates T-cell activation

Our observations suggest that coupling to membrane domains underlies the formation and location of protein condensates. Condensates have been previously implicated in T-cell signaling and activation (*18, 21*). Therefore, we hypothesized that coupling between domains and condensates might be important in T-cell function. We tested this hypothesis by uncoupling domains from condensates via two mutations of their critical linker LAT: (1) replacing the native LAT TMD with non-raft TMD (22 Leu)(*26, 27*)(Table S1) to create ‘nonraft LAT’ that still interacts with Grb2 but does not partition to raft domains or (2) mutating LAT’s 3 Tyr residues to Ala to create non-pY LAT, which can partition to rafts but cannot interact with Grb2. Fluorescence-tagged versions of these mutants (or wild-type LAT) were stably introduced into LAT-deficient Jurkat T-cells, and their capacity to facilitate T-cell activation was monitored by immunostaining against phosphorylated ERK (pERK) after plating on OKT3. JCam2.5 cells stably re-expressing wt-LAT showed strong pERK staining compared to the LAT-deficient negative controls (selected negative cells marked by asterisks in Fig 4I). In contrast, neither nonraft-LAT nor non-pY LAT showed pERK activation above LAT-deficient cells (Fig 4I and S22). Consistently, LAT-containing condensates were only observed in wt-LAT cells (Fig 4I, right). Cells expressing wt-LAT had ∼4-fold higher pERK signal than those expressing nonraft-LAT (Fig 4J). Importantly, these results were independent of PM LAT expression (Fig 4K) and even LAT phosphorylation (Fig S23), all of which were similar between wt-LAT and nonraft-LAT expressing cells. Thus, we conclude that coupling of membrane domains and cytoplasmic condensates via LAT is essential for activating signaling downstream of TCR ligation.

Synthesizing these observations, we find strong coupling of cytoplasmic protein condensates with lateral membrane domains in reconstituted models and in living Jurkat T-cells (Fig 4H). LAT condensates recruit specific lipids and proteins to their adjacent membrane by interactions between the LAT transmembrane domain and raft components (*26, 27*). LAT condensation stabilizes microscopic membrane domains (Fig 1E, 3H-L, 4A-B, S16, S19, 4H left), consistent with prevailing models of mammalian PMs containing dynamic nanodomains poised for coalescence by external inputs (*60*). Correspondingly, raft domains can nucleate and potentiate cytoplasmic condensates (Fig 1G, 3G, 4H right). Uncoupling LAT from rafts abrogates condensation and ERK activation downstream of TCR ligation. Therefore, we conclude that protein phase separation is thermodynamically and mechanistically coupled to lateral phase separation of membrane lipids to regulate the functional organization underlying immune cell signal transduction.

## Acknowledgements

Funding for IL was provided by the NIH/National Institute of General Medical Sciences (R35 GM134949, R01 GM124072, R21 AI146880), the Volkswagen Foundation (93091), and the Human Frontiers Science Program (RGP0059/2019). Funding for KRL was provided by NIH/National Institute of General Medical Sciences (R01 GM120351). Funding for MKR was provided by the Howard Hughes Medical Institute and the Welch Foundation (I-1544). JAD acknowledges support from The Hospital for Sick Children Research Institute. We acknowledge the labs of Sarah Veatch, Erdinc Sezgin, Xiaolei Su, Lawrence Samelson, Vasanthi Jayaraman, Jeanne Stachowiak, and Xiaodong Cheng for generous sharing of reagents, expertise, and/or equipment essential to this project.

