## Supplementary Figures and Methods for "Coupling of protein condensates to ordered lipid domains determines functional membrane organization"

**Supplementary materials for:** Coupling of protein condensates to ordered lipid domains determines functional membrane organization

Hong-Yin Wang, Sze Ham Chan, Simli Dey, Ivan Castello-Serrano, Jonathan Ditlev, Michael Rosen, Kandice R Levental*, Ilya Levental*

This file includes:
Figs. S1 to S23
Table S1
Movies S1 to S4
Supplementary Materials and Methods

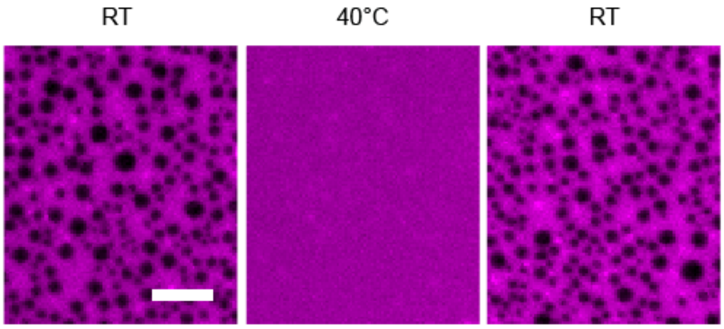


**Fig S1. Temperature reversibility of lipid domains on multi-bilayer membrane prepared through spin-coating.** Lipid composition: 31% DOPC, 33% DPPC, 33% chol, 2% DOGS-NTA, 0.2% trDHPE (Texas red-DHPE, Ld marker). Domains melt at high temperatures then reappear randomly when temperature is reduced. Scale bar is 15 µm.

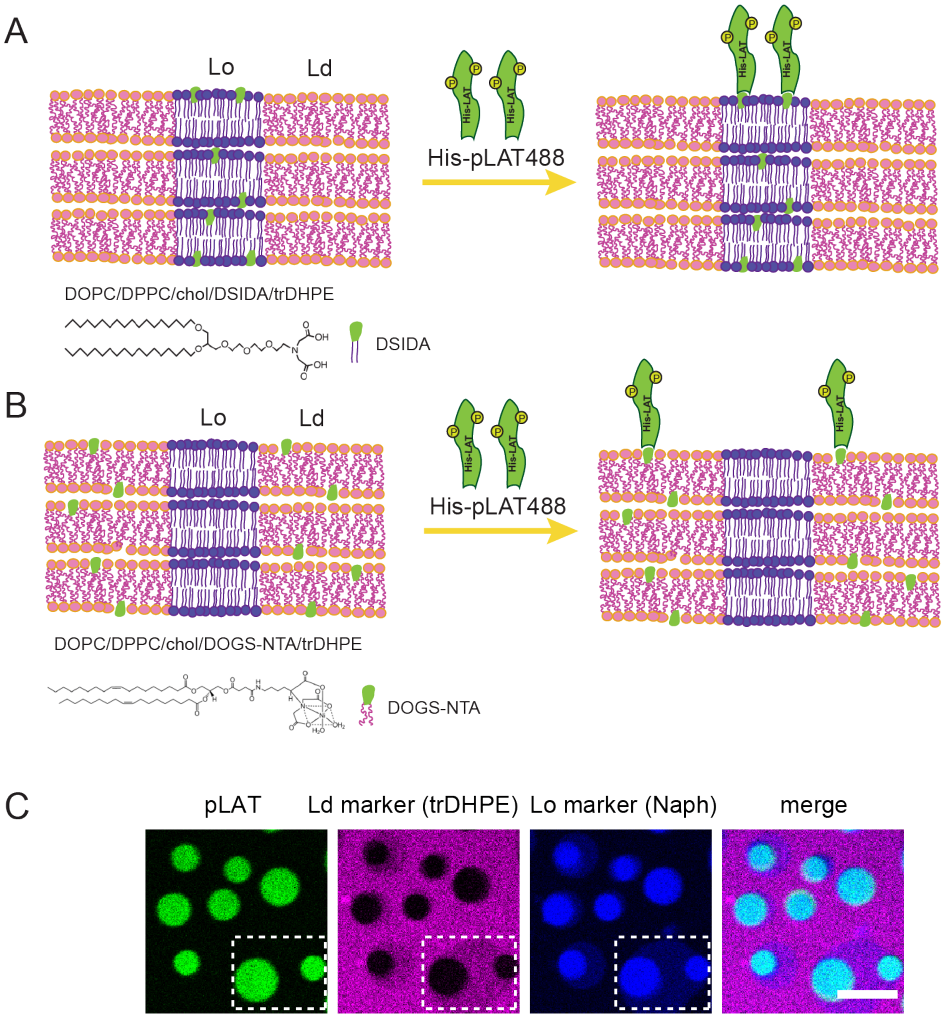


**Fig S2. Schematic of multi-bilayer experiment.** (A) Phosphorylated intracellular domain of LAT (His-pLAT488) is conjugated to Lo domain on multi-bilayer membrane through DSIDA lipids which contain saturated acyl chains (18:0). In some experiments, DPGS-NTA(Ni) lipids (DP-NTA; 16:0) were also used for the same purpose. (B) His-pLAT488 is conjugated to Ld domain through DO-NTA lipids. (C) Confocal images of configuration described in (A) with pLAT488 (green) conjugated to Lo domains (naphthopyrene, blue) through DSIDA (magenta = Ld marker, TR-DHPE). The region in the white rectangular box demonstrates an example of imperfect registration between the multiple bilayers: the top-most bilayer has two small circular Lo domains, which are overlying a larger, elliptical Lo domain in a lower bilayer. Evidence for this interpretation: (1) only domains on the top-most bilayer are accessible to LAT binding, so only domains on the top layer are green; (2) the region between the two circular domains has intermediate magenta and blue intensity, suggesting one type of domain overlying a different type (in this case, an Ld region overlying an Lo region in a lower bilayer). In general, most domains are registered across the bilayers, but some areas of incomplete registration are often observed. Scale bar is 5 µm.

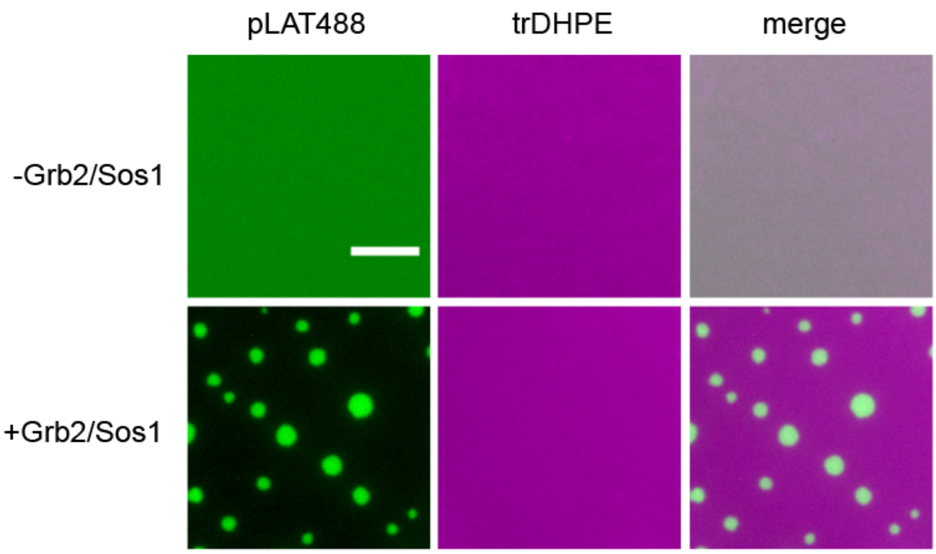


**Fig S3. LAT condensates form on a DOPC membrane after adding Grb2/Sos1.** Lipid composition: 98%DOPC, 2%DOGS-NTA(Ni), 0.2% trDHPE. Scale bar is 5 µm.

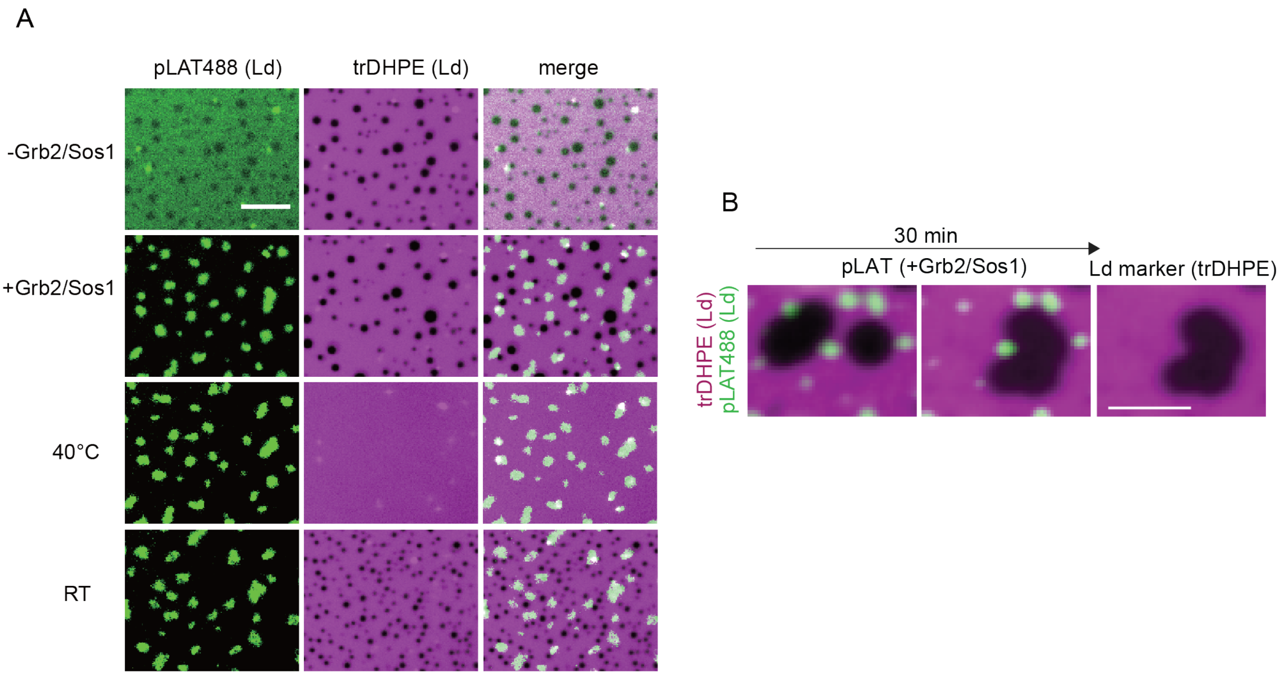


**Fig S4. LAT condensates conjugated to membrane through DOGS-NTA are excluded from Lo domains.** (A) *top* In the absence of Grb2+Sos1, pLAT coupled to the membrane via DOGS-NTA is dispersed throughout the Ld phase (48%DOPC, 32%DPPC, 18% chol, 2%DOGS-NiNTA, 0.2% trDHPE). (*2nd row*) Addition of Grb2+Sos1 leads to LAT coalescence into condensates, which form exclusively in the Ld phase. (*3rd row*) Membrane domains can be melted by raising the temperature to 40°C, while condensates persist. (bottom) Cooling below $T_{misc}$ leads to reappearance of Lo domains in new locations, with Lo domains formed upon cooling being absolutely prevented from forming underneath condensates. Scale bar is 5 µm (B) In this configuration Lo domains reappear exclusively away from the condensate-rich regions. This effect was particularly striking when protein condensates were present between two fusing membrane domains, where their presence defined the shape and boundary of newly fused domains. Scale bar is 5 µm.

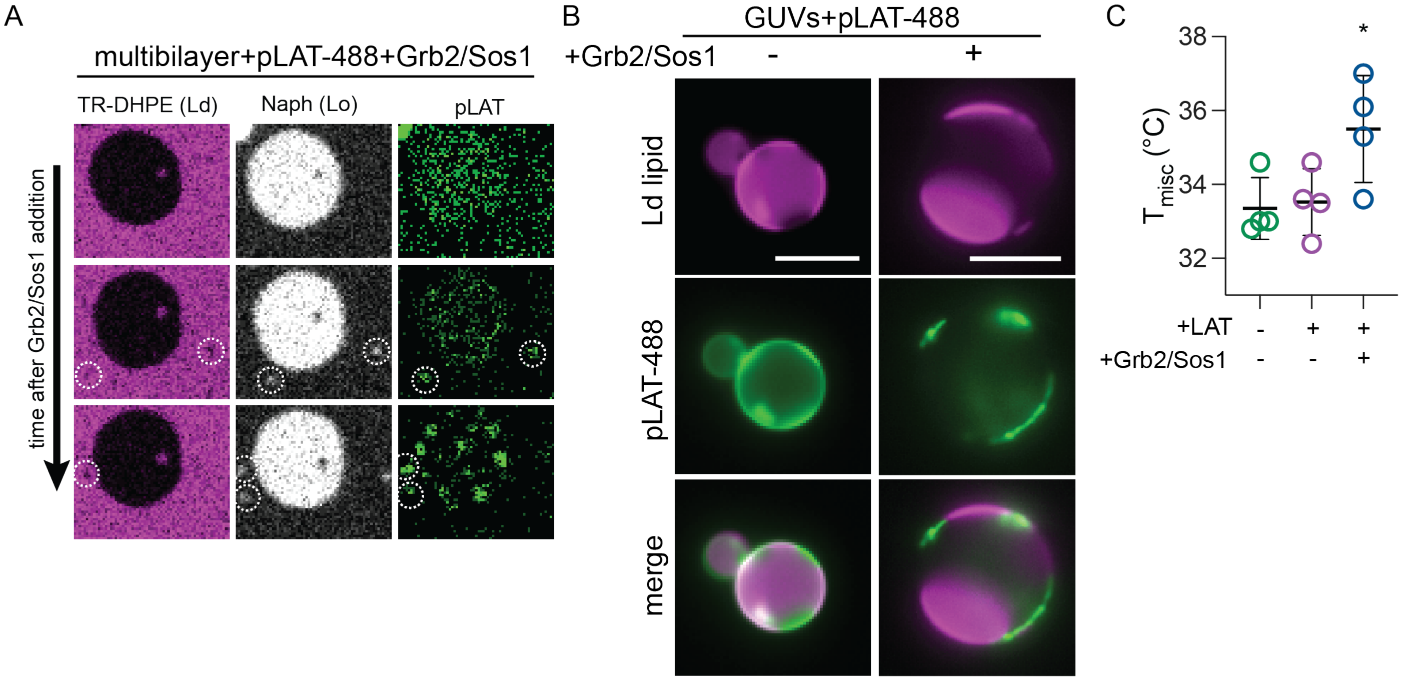


**Fig S5. LAT condensates conjugated to membrane with more biomimetic partitioning induce condensates in Lo domains and increase $T_{misc}$.** (A) In a phase-separating multi-bilayer (47% DPPC, 31% DOPC, 18% cholesterol, 2% DP-NTA, 1% DO -NTA, 1% Naphthopyrene, 0.2% trDHPE) where pLAT is ~2-fold concentrated in the Lo phase (mean $K_{p,Lo}$ = 2.1 ± 0.4) shown via naphthopyrene, white, middle panel), Grb2+Sos1 induced condensates in the Lo phase. Even when a nascent condensate forms in a region where the Ld phase dominates (white circles in the magenta bulk phase in the left column), small Lo domains (marked by naphthopyrene) are induced underneath the condensates. (B) Representative images of GUVs (38% DOPC, 36% DPPC, 20% chol, 4% DP-NTA, 2% DO-NTA, 0.04% LR-DHPE) with pLAT bound to both Lo and Ld phases with a mean $K_{p,raft}$=1.1±0.2. Shown are images taken at 33°C with GUVs in the absence and presence of Grb2+Sos1. (C) $T_{misc}$ in these GUVs was increased significantly by addition of 50 nM Grb2 / 25 nM Sos1. Shown is mean ± SD of the calculated $T_{misc}$ for 4 independent experiments. Details in Materials and Methods.

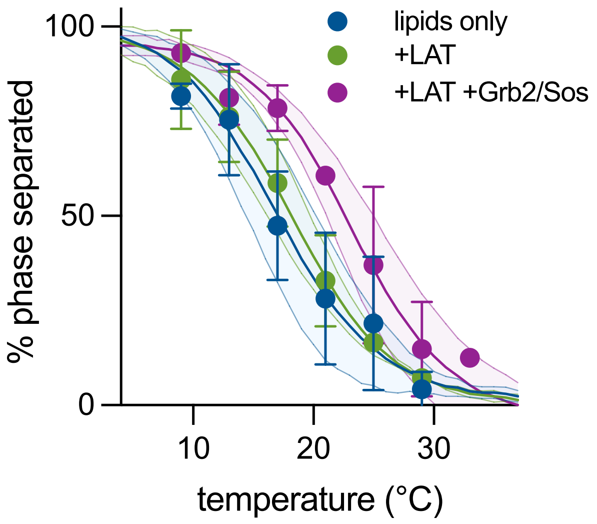


**Fig S6. Protein condensates enhance membrane phase separation.** Fraction of phase-separated GUVs is increased at higher temperatures with addition of Grb2/Sos1. Data points are mean ± SD from three independent experiments. The lines represent the average sigmoidal fits to the three independent experiments, with shading representing 95% confidence intervals on the fits.

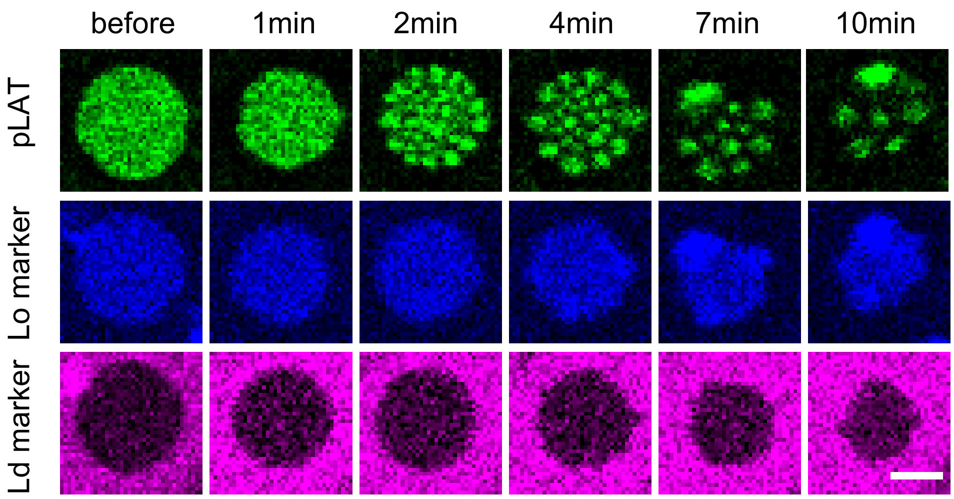


**Fig S7. Coarsening and arrangement of condensates inside Lo domain.** Confocal images of pLAT (green), the Lo marker naphthopyrene (blue), and Ld marker TR-DHPE (magenta) showing formation and coarsening of condensates inside the Lo domain after addition of Grb2/Sos1. The distribution of the condensates suggests that they are mutually repulsive.

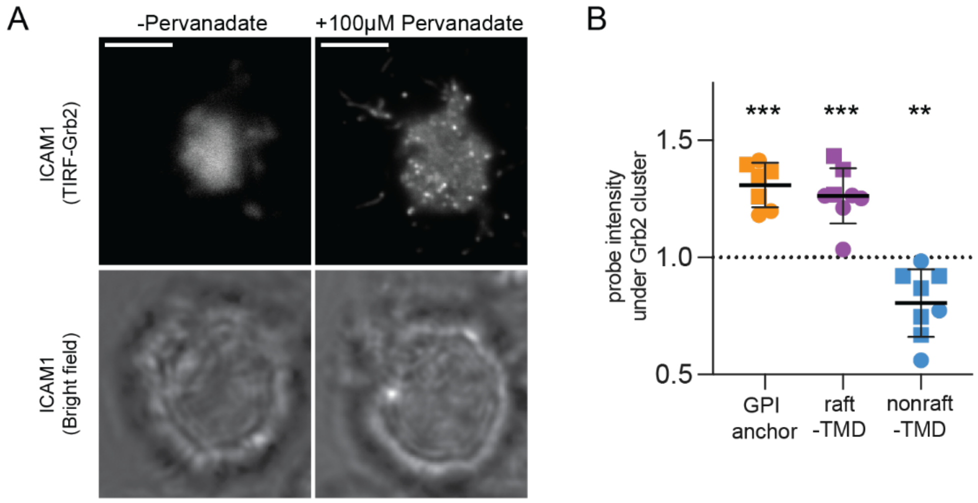


**Fig S8. Condensates formed in the absence of TCR ligation/clustering enrich for raft-prefering proteins.** (A) TIRF microscopy (top) and brightfield (bottom) images of Jurkats expressing Grb2-Scarlet plated on ICAM1. Treated with 100 µM pervanadate induced Grb2 condensates without TCR ligation, presumably by inhibiting dephosphorylation of LAT. (B) Quantification of relative enrichment of genetically encoded raft probes. Each point represents the mean enrichment (>1) or depletion (<1) of probes directly under Grb2 condensates relative to adjacent regions for individual cells across 3 independent experiments. Each cell included >10 Grb2 condensates. ***p<0.001, **p<0.01 for difference from 1 (no enrichment/depletion) of means of individual cells.

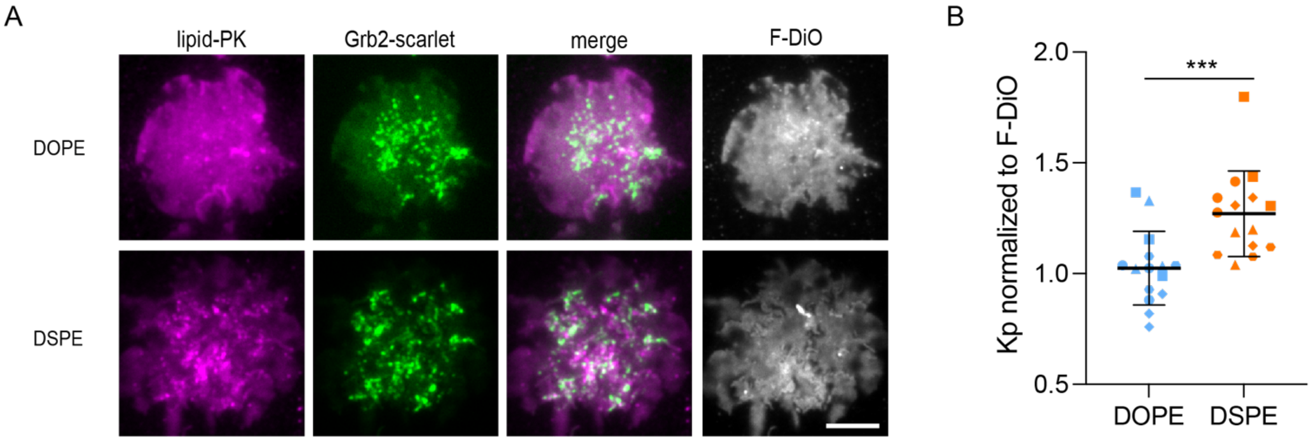


**Fig S9. Recruitment of raft-preferring lipids to Grb2 condensates in live cells.** Selectivity of Grb2 condensates for ordered membrane regions by imaging synthetic lipid fluorophores confined to the PM outer leaflet. (A) DSPE-PEG-KK114 consists of a PEG-linked far-red fluorophore coupled to a saturated lipid, imparting strong preference for ordered membranes (Honigmann, Mueller et al. 2013). TIRF imaging shows the recruitment of DSPE-PEG-KK114 (bottom row, magenta; see Table S1 for raft affinity quantification) to Grb2 condensates (green) in OKT3-activated Jurkat T-cells. In contrast, the non-raft-preferring lipid probe DOPE-PEG-KK114 did not show recruitment to Grb2 condensates (top row). The lipid dye F-DiO was used as internal control for both cases. Scale bar is 5 μm. (B) The partitioning of these lipid probes to condensate-associated membranes was quantified as a normalized partition coefficient (Kp) relative to the lipid dye (F-DiO). Kp is calculated as the fluorescence intensity of dye in condensate-associated membrane divided by its intensity outside condensate regions. This Kp value is then normalized to the Kp of F-DiO. Points represent individual condensates and shapes represent individual cells. ***$p<0.001$ from student's t-test.

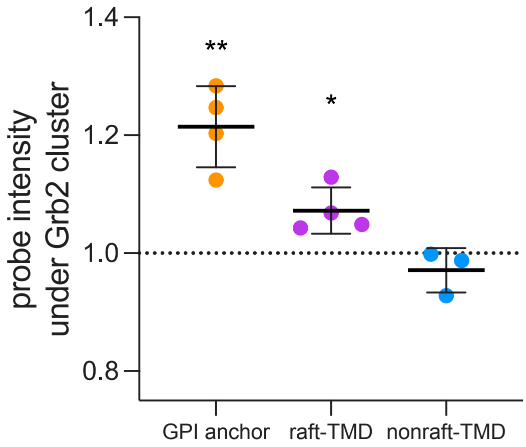


**Fig S10. Grb2 condensates enriched for raft-preferring probes when imaged at 37°C.** Quantification of enrichment of genetically encoded raft partitioning probes in membrane associated with Grb2 condensates formed in OKT3-activated Jurkat cells and imaged at 37°C. Each point represents the mean enrichment (>1) or depletion (<1) of probes directly under Grb2 condensates relative to the adjacent region for individual cells across 3 independent experiments. Each cell included >10 Grb2 condensates. **$p<0.01$, *$p<0.05$ for difference from 1 (no enrichment/depletion) of means of individual cells.

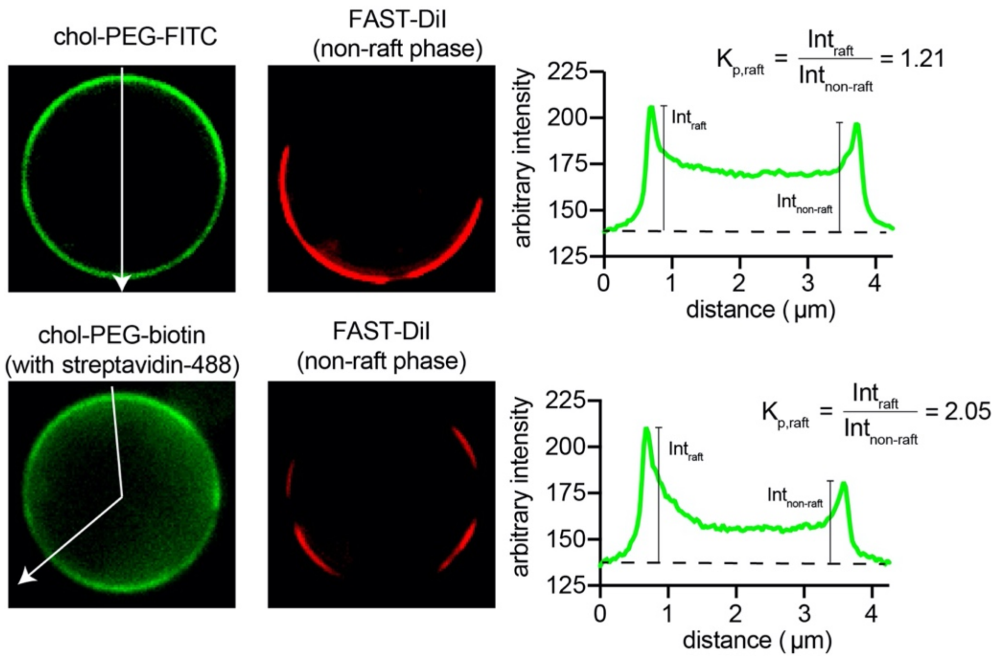


**Fig S11. Example of raft affinity quantification in GPMVs.** Chol-PEG-FITC (green) is slightly enriched in raft phase of phase separate GPMVs. The line scan along the white line shows that (background subtracted) FITC intensity is ~20% higher in raft phase, compared to non-raft phase marked by FAST-DiI (red). Labeling and crosslinking cholesterol-PEG-biotin (chol-PB) with fluorescent avidin (Av488) increased the raft phase affinity of crosslinked chol-PB.

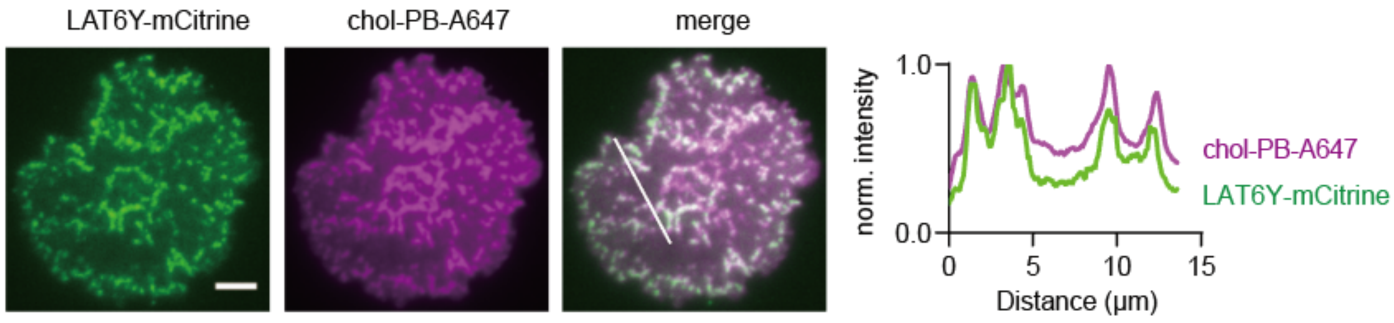


**Fig S12. Colocalization of LAT with cholesterol-rich domains in live activated T cells.** TIRF imaging of colocalization of LAT condensates with chol-rich domains formed on an activated Jurkat T-cell. The cell line used is a LAT-deficient line (Jcam2.5) stably expressing a LAT variant containing six tyrosine residues for Grb2 binding. Scale bar is 5 μm.

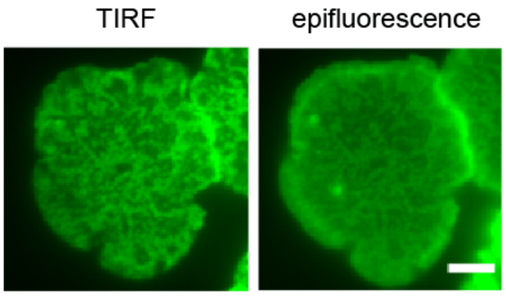


**Fig S13. Both TIRF (left) and epifluorescence (right) reveal chol-rich domains in activated, chol-PB-labeled Jurkats.** Chol-PB visualized by staining with fluorescent streptavidin. Scale bar is 5 μm.

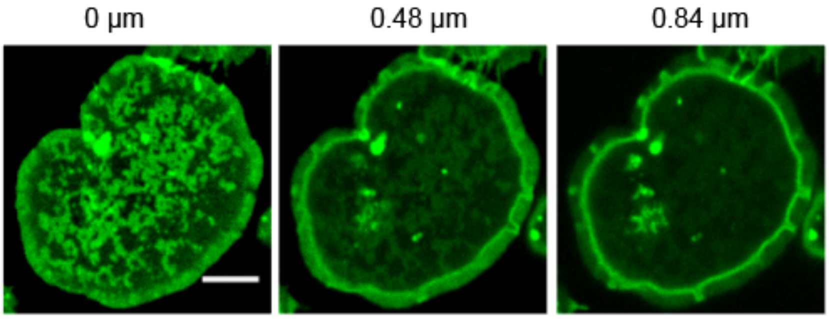


**Fig S14 Confocal imaging of different focal planes of chol-rich domains formed on activated Jurkat T-cell membranes.** Imaging shows that chol-rich domains are only observed at the surface of the cells and are not microscopically resolvable membrane invaginations. Scale bar is 5 µm.

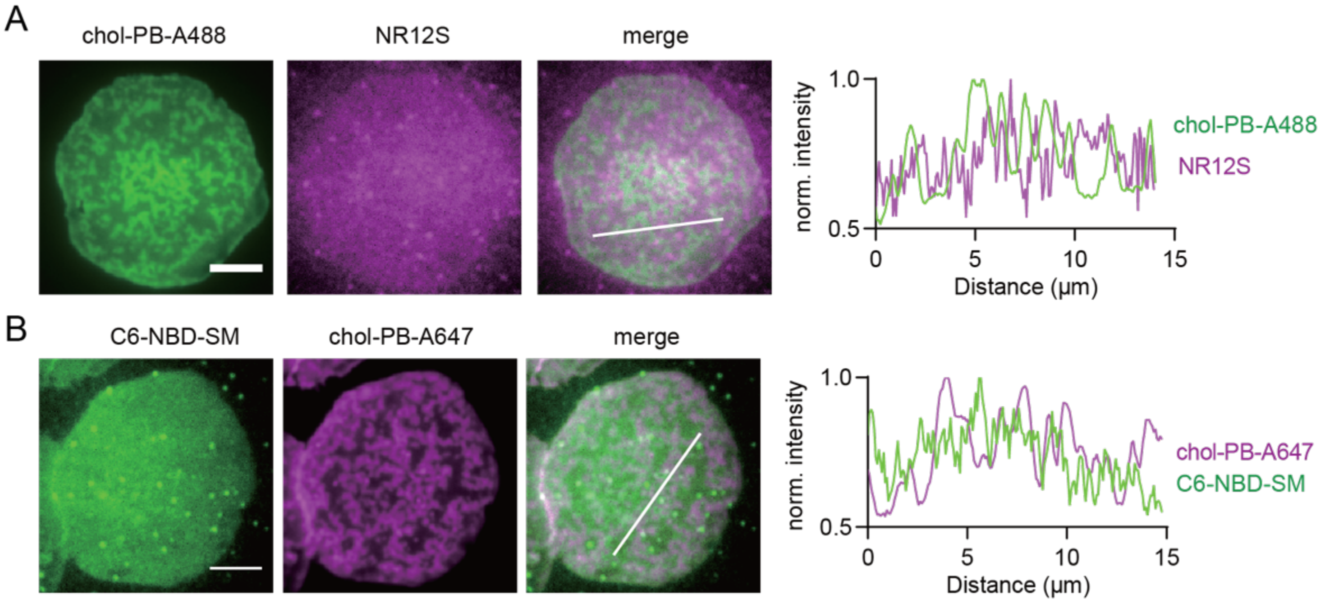


**Fig S15. TIRF imaging of chol-rich domains formed on activated Jurkat T-cells.** Before activation, Jurkat T-cells were labeled with chol-PB-A488 and NR12S. After activation, neither NR12S (A) nor C6-NBD-SM (B) is co-enriched with chol-PB-A488, indicating the chol-rich domains are not membrane topography features, like membrane invaginations or deformations. Scale bar = 5 µm.

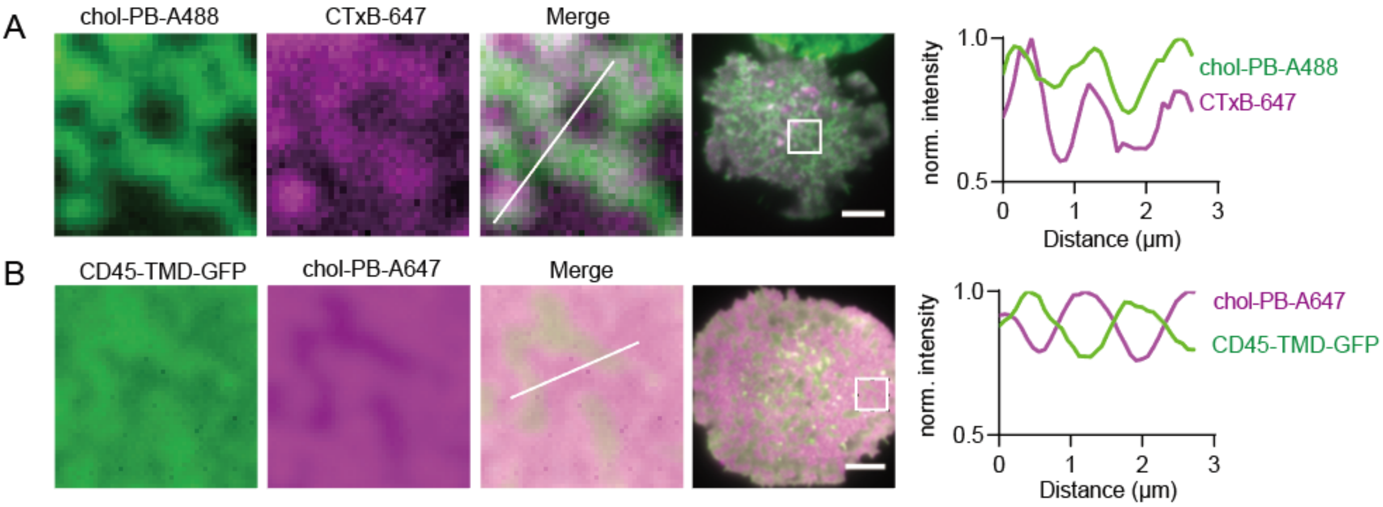


**Fig S16. Stable cholesterol-rich domains (crosslinked chol-PB) formed in activated T-cells enrich in raft markers and exclude nonraft proteins.** (A) TIRF imaging of chol-PB-A488 overlying with raft marker CTxB. Plot shows relative fluorescence intensity along the line trace shown in white. (B) TIRF imaging shows exclusion of nonraft CD45-TMD (Table S1) from chol-rich domains (chol-PB-A647). Scale bars are 5 μm.

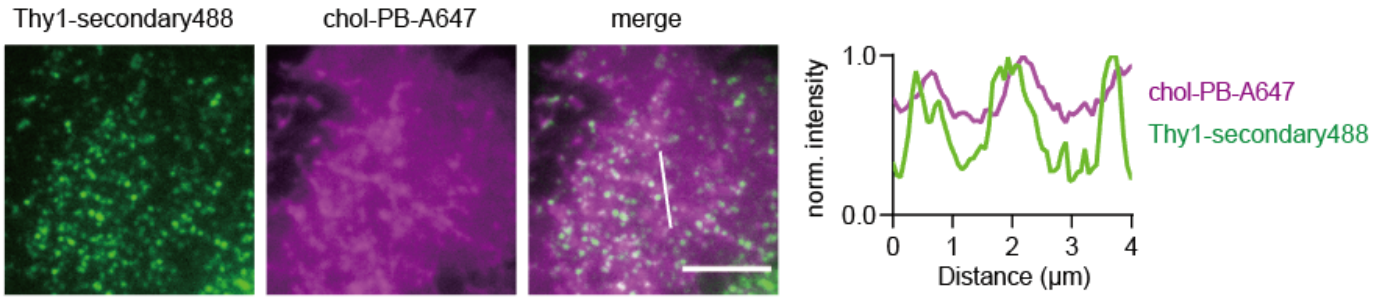


**Fig S17.** TIRF imaging shows the overlapping of GPI-AP clusters (Thy1 crosslinked by secondary antibodies) with cholesterol-rich domains labeled with chol-PB-A647 on the plasma membrane of an activated Jurkat T-cell. Scale bar is 5 μm.

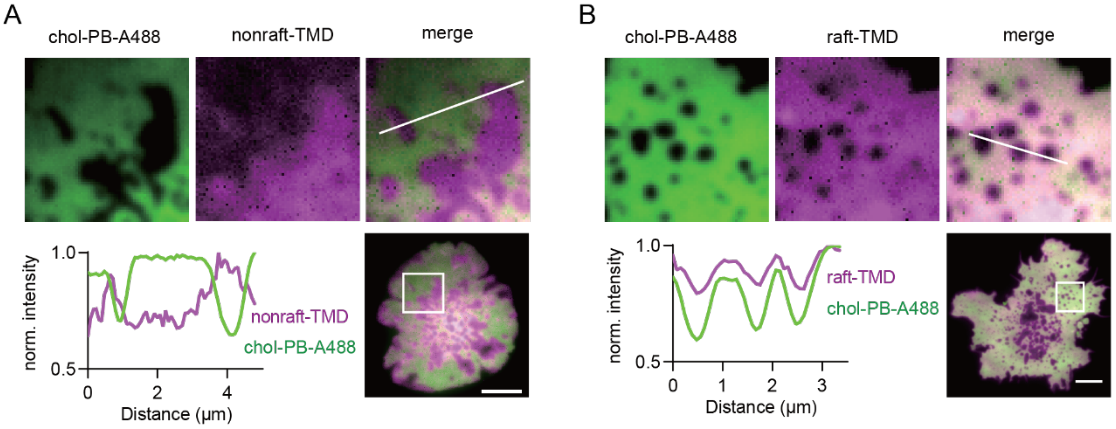


**Fig S18. TIRF imaging of exclusion of nonraft-TMD and recruitment of raft-TMD by chol-rich domains formed in Jurkat T-cells activated on supported lipid bilayers (SLB).** SLB were prepared through vesicle fusion with a lipid composition of 98%DOPC, 1% DSPE-PEG2k-biotin, 1% DOGS-NTA(Ni). His-tagged ICAM-1 was linked through DOGS-NTA(Ni) lipids to enhance cell attachment on SLB. Biotin-OKT3 was linked on the SLB through streptavidin and DSPE-PEG2k-biotin lipids. Chol-rich domains are depleted of nonraft-TMD (A) and enriched in raft-TMD (B) probes. For detailed methodological information, please refer to the supplementary materials. Scale bars are 5 μm.

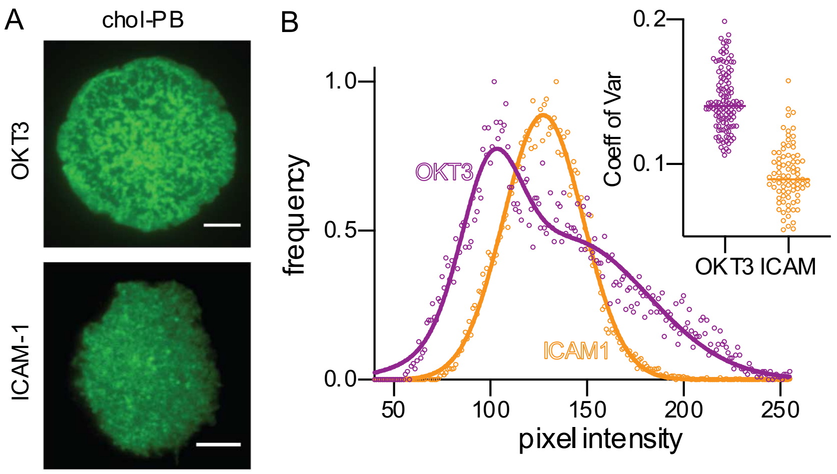


**Fig S19. Micron-scale chol-rich domains are only observed in the presence of LAT condensates (ie on OKT3).** (A) ICAM1-coated coverslips (bottom), which promote robust Jurkat cell attachment without TCR-mediated signaling activation, do not induce micron-scale chol-rich domains as OKT3 does (top). Scale bars = 5 µm. (B) Chol-rich versus chol-poor domains appear as bi-modal distribution of pixel intensities. (inset) The spread of pixel intensities is quantified via coefficient of variation (st. dev. / mean) which is significantly larger in activated cells with condensates (OKT3) compared to adhered cells without condensates (ICAM). Data points represent 5x5 µm regions from 5-10 cells/condition; ***$p<0.001$ for differences between cells.

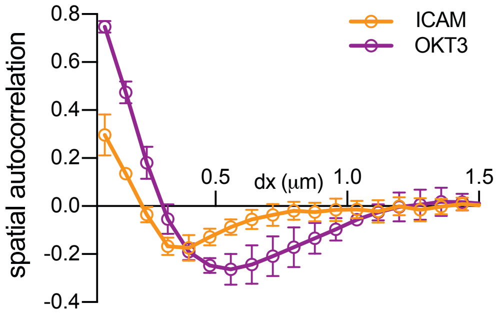


**Fig S20. Spatial autocorrelation of pixel intensities reveals greater magnitude and length scale of chol-rich domains on OKT3 than ICAM1.**

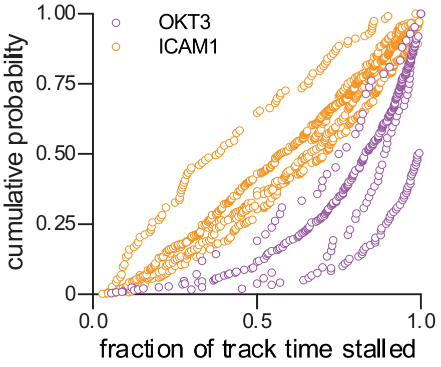


**Fig S21. Stalled fractions of individual tracks shown as cumulative distribution on a per-cell basis.** Thy1 clusters tracked over time in TIRF time-lapse movies. The Thy1 cluster tracks on OKT3 are generally stalled for the majority of the tracks as compared to the Thy1 clusters on ICAM1.

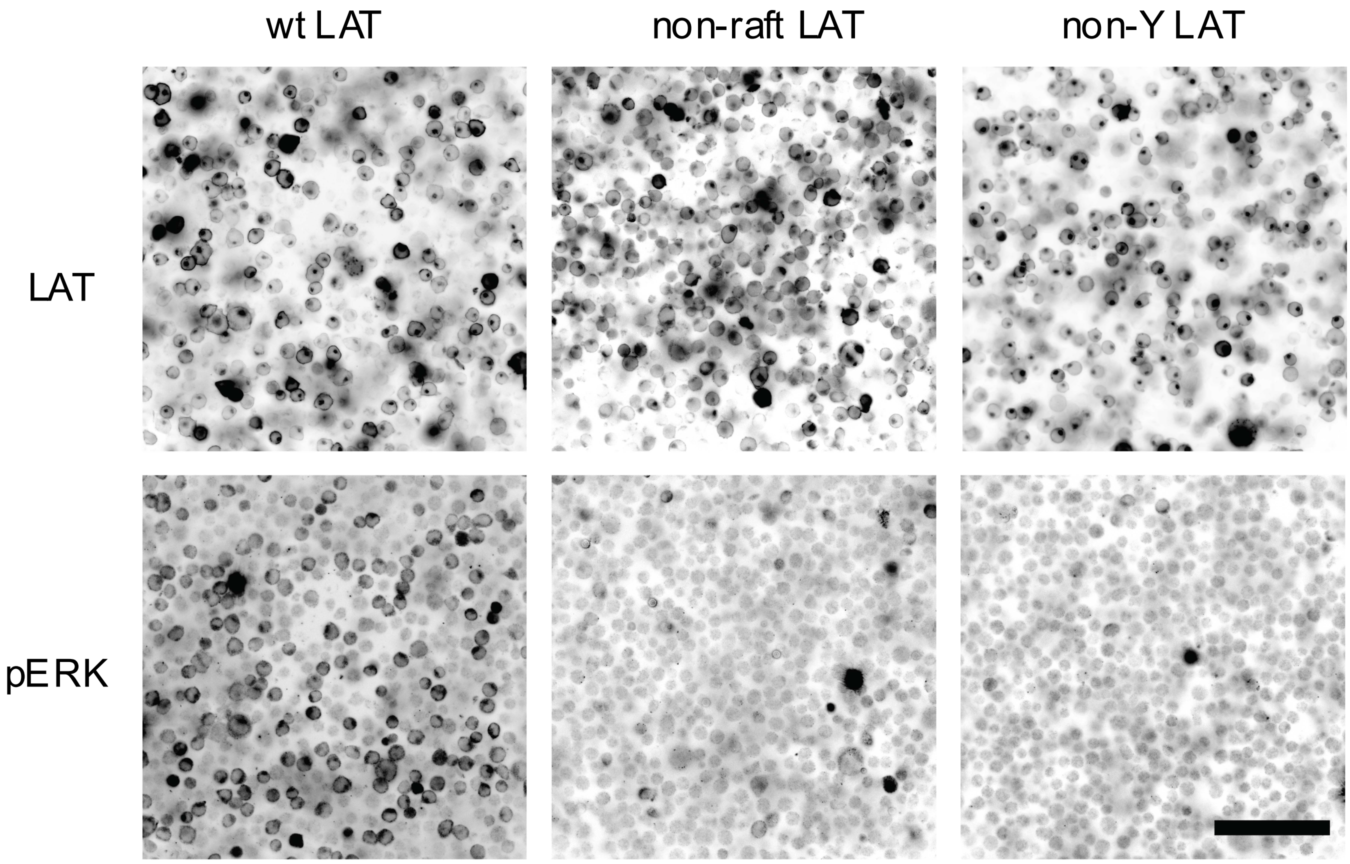


**Fig S22. Only cells containing WT LAT (i.e. containing both the raft-targeting TMD and phosphorylatable Tyr to form condensates) can activate pERK on OKT3.** Stable cell lines expressing either mCitrine-tagged wt-LAT, non-raft LAT, or non-pY LAT were mixed 1:1 with LAT-deficient Jcam2.5 cells (not fluorescent) and plated for 10 min at RT on OKT3-coated coverslips. IF staining of pERK (bottom panel) showed that only wt-LAT-positive cells showed strong pERK signal, while neither non-raft nor non-pY LAT cells showed pERK staining above the background levels of LAT-deficient Jcam2.5 cells. Scale bar is 100 µm.

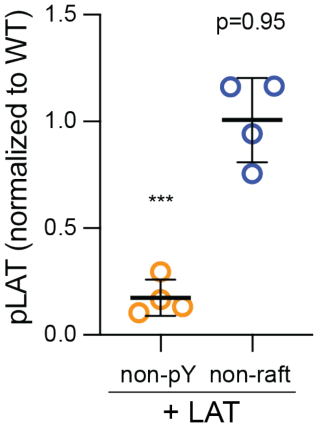


**Fig S23. Phosphorylation of LAT is not affected by its raft affinity.** Immunofluorescence of JCam2.5 LAT-deficient cells transfected with GFP-tagged wild-type LAT, non-raft LAT (i.e. with allL-TMD), or non-pY LAT (phosphorylation-null LAT: Y171F, Y191F, and Y226F). The non-pY LAT serves as the pLAT/LAT negative control. The mean pY-LAT / LAT ratio was calculated for at least 25 cells over 4 independent experiments. The means of each experiment were calculated and normalized to WT LAT. Whereas the non-phosphorylatable LAT is significantly lower than WT, both raft (WT-) and nonraft (allL-) LAT are phosphorylated to similar extent, suggesting that nonraft-LAT's inability to form condensates is not due to lack of phosphorylation (i.e. upstream signaling). Mean ± SD of individual experiments are shown. ***$p<0.001$; one-sample t-test compared to theoretical mean of 1.

**Supplementary Table S1.** Raft phase affinity of various probes used in this study, quantified in Giant Plasma Membrane Vesicles (see Fig S11 and methods for details).

| Probe | $K_{p,raft}$ (Mean ± SD) |
|---|---|
| raft-TMD (i.e. LAT TMD tagged with RFP) | 1.21 ± 0.14 |
| nonraft-TMD (i.e. allLeu TMD tagged with RFP) | 0.37 ± 0.10 |
| CD45-TMD | 0.49 ± 0.12 |
| cholesterol-PEG-FITC | 1.26 ± 0.05 |
| cholesterol-PEG-biotin (labeled with streptavidin) | 1.87 ± 0.29 |
| DOPE-PEG-KK114 | 0.79 ± 0.07 |
| DSPE-PEG-KK114 | 1.37 ± 0.18 |
| monomeric GPI-AP (GPI-GFP) | 1.57 ± 0.15 |
| dimeric GPI-AP (Thy1 with primary antibody) | 2.10 ± 0.03 |
| GPI-AP cluster (Thy1 with secondary antibody) | not quantifiable, essentially all clusters in Lo |

**Supplementary Movie 1.** TIRF movie of secondary-antibody crosslinked Thy1 clusters (magenta) diffusing on the surface of an activated Jurkat cell, with Grb2-mScarlet (green) marking membrane-associated protein condensate. Thy1 clusters stops diffusing underneath the Grb2 condensate. Frame size = 2x2 μm ; time step = 120 msec.

**Supplementary Movie 2.** TIRF movie of an immobilized secondary-antibody crosslinked Thy1 cluster (magenta) on the surface of an activated Jurkat cell. Initially, other diffusing clusters are recruited and immobilized, then a Grb2-mScarlet protein condensate (green) forms. Frame size = 2x2 μm ; time step = 120 msec.

**Supplementary Movie 3.** TIRF movie of secondary-antibody crosslinked Thy1 clusters (green) diffusing on the surface of an activated Jurkat cell, with Grb2-mScarlet (magenta) marking membrane-associated protein condensate. Two large condensates recruit and immobilize Thy1 clusters. Other clusters immobilize in nearby regions of the membrane without condensates. Frame size = 2x2 μm, time step = 70 msec.

**Supplementary Movie 4.** TIRF movie of secondary-antibody crosslinked Thy1 clusters (green) diffusing on the surface of an activated Jurkat cell, with Grb2-mScarlet (magenta) marking membrane-associated protein condensate. Most immobilized Thy1 clusters are localized underneath Grb2 condensates. Some clusters are immobilized in other regions of the membrane. Newly immobilized Thy1 cluster (frame 52, middle of frame) nucleates subsequent condensate formation. Then newly formed condensate (frame 125, top right of frame) recruits and immobilizes a Thy1 cluster. Scale bar = 1 μm, time step = 70 msec.

**MATERIALS AND METHODS:**

*Materials:* Cholesterol-PB and cholesterol-PEG-FITC were purchased from Xi'an Ruixi Biological Tech). The following reagents were purchased from BioLegend: anti-Thy1 primary (1°Ab) antibody (mouse anti-human and mouse anti-rat, both unlabeled, FITC-labeled, and AlexaFluor 647-labeled), OKT3 (anti-CD3 antibody). ICAM1 was purchased from Sino Biological. The following were purchased from Thermo Scientific: phospho-Erk1/2 (Tyr204) Fast-DiO, Fast-DiI, DiD, Streptavidin-AlexaFluor 488 or 647. DSPE-PEG-KK114 and DOPE-PEG-KK114 were a gift from Dr. Erdinc Sezgin's lab. DOPC, DPPC, cholesterol, trDHPE, LissRhod-DOPE, DOGS-NTA(Ni), DPGS-NTA (Ni) and C6-NBD-SM were purchased from Avanti Polar Lipids. DSIDA was a gift from Dr Jeanne Stachowiak's lab. NR12S was kindly provided by Andrey Klymchenko. Naphtho[2,3-a]pyrene was purchased from TCI America. Myriocin was purchased from Cayman Chemials. Zaragozic Acid was purchased from Sigma.

*Purified proteins:* Human Grb2 (aa 1-217), Sos1 (aa 1117-1319), and LAT (aa 48-233) with eight tyrosine phosphorylation sites were purified from bacterial expression via GST-tags, as previously described (Su, Ditlev et al. 2016). pLAT8Y-alexa488 (henceforth referred to as pLAT) was created by phosphorylation with GST-ZAP70 and labeled by maleimide-conjugated dye, as described (Su, Ditlev et al. 2016). Concentrations used for reconstitution experiments were previously established as being within the physiological range (Su, Ditlev et al. 2016).

*Planar lipid multi-bilayers prepared by spin-coating:* supported lipid multi-bilayers were prepared according to published protocols (Zeno, Johnson et al. 2016). Briefly, lipid solutions were mixed in chloroform and dried under $N_2$ gas stream. Films were then dissolved to 3 mg/mL spin-coating solution of 97% hexane and 3% methanol. A freshly cleaved mica substrate (25mm x 25 mm, G1, Ted Pella Inc, #56-25) was placed on a custom-built spin coater and 60 µl coating solution was added at 3000 rpm and spun for 30 seconds. The mica coated with lipid film was placed under vacuum for >2 h to completely evaporate the organic solution. Next, a microfluidic chamber (5 µL volume) was constructed with the coated mica substrate as its base. Inlet tubing with a small diameter was used to reduce the dead volume to 1.4 μL/10 cm. To achieve uniform rapid hydration, 30 μL of hydration buffer (150 mM NaCl, 20 mM Tris, pH7.4) were pumped into the chamber in <2 seconds. The resulting supported multi-bilayer stack was then allowed to equilibrate at room temperature for 10 min prior to addition of proteins. For membranes with ternary lipid compositions required for phase separation, the multi-bilayer stack was heated to 45°C on a Peltier stage (Warner Instruments) to melt domains and fully equilibrate the bilayer. The whole system was then equilibrated for 10 min at each measurement temperature.

*Observations of protein condensates on lipid bilayer membranes:* to maintain integrity of supported membranes in the microfluidic chamber, all solution changes were done through a syringe pump with a flow rate of 1 μL/min. The chamber was first blocked in hydration buffer with 0.1% BSA (blocking buffer) for 20 min at 23°C. All subsequent solutions were in blocking buffer. First, His-tagged pLAT (100 nM) was added to the chamber and incubated for 20 min (for membrane with DSIDA lipids, 200 μM $CuCl_2$ was included in blocking buffer). Unbound pLAT was washed out by pumping with blocking buffer for 10 min. Grb2 (1.3 μM) and Sos1 (80 nM) were premixed and pumped into the chamber, then condensate formation was monitored microscopically. Temperature was controlled by a Peltier stage (Warner Instruments).

*Lipid phase separation induced condensate formation:* To lower the concentration of DSIDA and LAT to a regime where no condensates are observable on a uniform DOPC membrane, but condensates can be formed in phase separated membranes, experiments were carried out as mentioned above with some modifications. In detail, membranes (lipid composition was listed below as Fig 1G) were prepared through spin-coating on a circular mica substrate (12 mm, G1, Ted Pella Inc, #50-12) which was cleaved and mounted on a coverslip (#1) through an UV-curing optical adhesive (Thor Labs, #NOA81). Disc-shaped solidified PDMS (polydimethylsiloxane) material (around 4 mm thickness and 15 mm diameter) with a 3 mm hole poked by biopsy punch (Electron Microscopy Sciences, #69038-03) was mounted on the spin-coated lipid films to form a coverless chamber. Prior to chamber assembly, to reduce nonspecific binding of proteins,

PDMS material was incubated with 1% BSA in hydration buffer at 4C overnight and washed 3 times. In the following experiment, only hydration buffer was used without any BSA and solution exchanging in the chamber was achieved through 10 ul volume exchanging for 5 times. After hydration with 35 ul hydration buffer, membrane were equilibrated at 55°C for 5 min and cooled down to RT for 30 min. 200 µM $CuCl_2$ was incubated with the membranes at RT for 10 min after which 20 nM pLAT was incubated with the membrane for 20min. After washing out the unbound pLAT, Grb2 (100 nM) and Sos1 (100 nM) were premixed and added into the chamber, then condensate formation was monitored microscopically. Same method was used for the sample of mixed DP-NTA and DO-NTA lipids.

*GUV preparation and $T_{misc}$ measurement:* GUVs were produced by electroformation as previously described (Levental, Lorent et al. 2016). Briefly, lipid solutions in chloroform (1.5 mg/mL) were applied to two electrodes of a custom-built chamber and dried under vacuum for 1 h. The electrodes were then submerged in a chamber preloaded with 400 µL sucrose (0.4 M). Alternating current (2 V, 10 Hz, sine wave) was applied to the electrodes for 1.5 h at 70°C. The GUVs were detached from the electrodes with a square wave current of 2 Hz, 2 V for 15 min. The GUV suspension was then cooled and diluted into 2 mL isotonic glucose-containing buffer (100 mM NaCl, 20mM Tris, pH 7.4, 0.1% BSA, 200 µM $Cu^{2+}$; 400 mOsm adjusted with glucose). To form condensates, 200 nM pLAT was incubated with the GUVs for 20 min at 23°C with gentle agitation. Then unbound pLAT was washed out by centrifugation twice with 100 rcf for 5 min. Then Grb2 (final concentration: 135nM) and Sos1 (final concentration: 67.5nM) were premixed and added to the GUV solution with gentle mixing. GUVs were imaged by epifluorescence microscopy. To measure the thermotropic phase behavior (i.e. $T_{misc}$), GUVs were added to a coverslip chamber and temperature was controlled by a microscope-mounted Peltier stage (Warner Instruments). The fraction of phase separated GUVs was measured as a function of temperature, producing characteristic sigmoidal curves (as in Fig S6) with $T_{misc}$ defined as the temperature at which 50% of GUVs were phase separated. The same method was used to measure the effect of crosslinking GPI anchors on $T_{misc}$ in GPMVs (Fig 4D).

*Compositions for supported bilayers and GUV experiments:*

Fig 1A: 31% DOPC, 27% DPPC, 40% cholesterol, 2% DSIDA, 0.04% trDHPE

Fig 1B-C: 50% DOPC, 30% DPPC, 18% cholesterol, 2% DSIDA, 0.2% trDHPE

Fig 1D, Fig S5A: 47% DPPC, 31% DOPC, 18% cholesterol, 2% DPGS-NTA, 1% DOGS -NTA, 1% Naphthopyrene, 0.2% trDHPE

Fig 1E-F, Fig S6: 28% DPPC, 30% DOPC, 40% cholesterol, 2% DSIDA, 0.04% LissRhod-DOPE

Fig 1G, left panel: 99.3% DOPC, 0.5% DSIDA, 0.2% trDHPE

Fig 1G, right panel, Fig S2C, Fig S7: 48.3% DOPC, 32% DPPC, 18% chol, 0.5% DSIDA, 0.2% trDHPE, and 1% naphthopyrene.

Fig S1: 31% DOPC, 33% DPPC, 33% chol, 2% DOGS-NTA, 0.2% trDHPE

Fig S3: 98%DOPC, 2%DOGS-NTA, 0.2% trDHPE

Fig S4: 48% DOPC, 32% DPPC, 18% cholesterol, 2% DOGS-NTA, 0.2% trDHPE

Fig S5B-C: 36% DPPC, 38% DOPC, 20% cholesterol, 4% DPGS-NTA, 2% DOGS-NTA, 0.04% LissRhod-DOPE

Fig S16 of SLB for Jurkat activation: 98%DOPC, 1% DSPE-PEG2k-biotin, 1% DOGS-NTA-Ni

*Plasmids, cell culture and transfection:* raft-TMD (tr-LAT), nonraft-TMD (tr-allL), and GPI-GFP were previously described (Levental, Lingwood et al. 2010, Lorent, Diaz-Rohrer et al. 2017). Grb2-GFP was purchased from Addgene (#86873). Grb2-mScarlet was a gift from Dr Lawrence Samelson's lab. CD45-TMD was a gift from Dr Sarah Veatch's lab. LAT-deficient Jcam2.5 cells and Jurkat T-cells that stably express LAT6Y-mCitrine and non-pY LAT-mCitrine were kindly provided by Xiaolei Su (Yale University) as previously described (Su, Ditlev et al. 2016). Jurkat T-cells (ATCC) were cultured in RPMI medium supplemented with 10% fetal bovine serum (FBS), 100 units/mL penicillin, and 100 µg/mL streptomycin. Rat basophilic leukemia (RBL-2H3) cells (ATCC) were grown in medium containing 60% modified Eagle's medium, 30% RPMI, 10% FBS, 100 units/mL penicillin, and 100 µg/mL streptomycin. Hela cells were cultured in DMEM medium supplemented with 10% FBS, 100 unites/mL penicillin, and 100 μg/mL streptomycin. Cells were transfected through electroporation (Nuclefector, Lonza).

*Stable cell lines of wt-LAT and non-raft LAT Jurkats:* LAT-deficient Jcam2.5 cells were transfected and expressed with either GFP-tagged LAT or GFP-tagged LAT with its TMD only being mutated to Leucine residues (allL; non-raft). 1 mg/ml G418 was added to the Jurkat medium, and cells were selected for 2 weeks. Finally, FACS was used to sort for for the 10% of cells with the greatest GFP signals, and the stable cell lines were obtained.

*Membrane probes and plasma membrane staining:* chol-PB and chol-PEG-FITC (10 μg/mL) were incubated with cells in PBS at 4°C for 5 min. For avidin crosslinking, 2 μg/mL streptavidin488 or 647 were incubated with chol-PB labeled cells at 4°C for another 5 min. For DSPE-PEG-KK114 and DOPE-PEG-KK114), 2 μg/mL of the probes were incubated with the cells at 37°C for 5 min. For C6-NBD-SM, 2 μg/mL was used to stain the cells at 4°C for 5 min. For NR12S, cells were stained with 30 nM at 4°C for 5 min. After membrane staining, the cells were washed with PBS and resuspended in imaging buffer (Tyrode's extracellular buffer: 25 mM HEPES, 150 mM NaCl, 5 mM KCl, 5.4 mM glucose, 1 mM $CaCl_2$, 0.4 mM $MgCl_2$, pH7.2).

*Labeling and crosslinking endogenous Thy1*: Jurkat T cells were first labeled with monoclonal anti-Thy1 primary (1°Ab) antibody, (5 μg/mL for unlabeled 1°Ab; 4 μg/mL for Alexa-647 or FITC-labeled 1°Ab) in 0.1% BSA containing PBS buffer (staining buffer) on ice for 15 min. For secondary antibody labeling, Alexa488/647-labeled donkey anti-mouse antibody (10 μg/mL) were used to stain the cells in staining buffer on ice for 15 min. After labeling, cells were washed once with staining buffer and resuspended in Tyrode's extracellular buffer.

*Jurkat T cell activation on OKT3-coated coverslips and TIRF imaging:* Glass-bottomed 4-well dishes (Matsunami, #D141400) were coated with OKT3 (anti-CD3 antibody) by incubating with 5 μg/mL OKT3 in PBS overnight at 23°C. After thoroughly washing with PBS, the coated coverslips were blocked with 5% BSA in PBS for 1 h at 23°C, then maintained with 100 μL imaging buffer (Tyrode's extracellular buffer: 135 mM NaCl, 5.0 mM KCl, 1.8 mM CaCl2, 1.0 mM MgCl2, 5.6 mM glucose, 20 mM HEPES, 1.0 mg/mL BSA, pH 7.4). Suspended Jurkat T-cells (labeled as above) were added to the coverslip chambers, activating as they made contact with the coated coverslip surface. Jurkat activation was monitored (at RT if not indicated, or 37°C where indicated) with TIRF microscopy with 100x TIRF objective (NA=1.4) at the TIR incident angle. Excitation laser wavelengths used were 488 nm, 561 nm, and 647 nm according to fluorophores and corresponding emission filters were applied to reduce crosstalk between different channels. For non-activating surfaces, BSA (5 μg/mL) or ICAM-1 (5 μg/mL) were incubated with coverslips under similar conditions as OKT3.

*Crosslinking Thy1 induced condensates formation:* For 1°Ab crosslinking experiment, Jurkat cells were directly added on anti-Thy1 primary (1°Ab) antibody (5 μg/ml) incubated coverslips. For 2°Ab crosslinking experiment, Jurkat or LAT-deficient Jcam2.5 cells were first labeled with primary Thy1 antibody as mentioned above and then added to coverslips that were previously coated with Alexa647-labeled donkey anti-mouse antibody (5 μg/mL, RT overnight). Condensate formation in these cells were observed through TIRF microscope as mentioned above.

*Immunostaining (IF) of phosphorylated ERK (pERK) and phosphorylated LAT (pLAT):* Jcam2.5 cells were mixed 1:1 in cell number with either wt-LAT, nonraft-LAT, or non-pY LAT Jurkat cells. For pLAT IF, 1 million cells of GFP-labeled wt-LAT, non-raft LAT and non-pY LAT cells were used in each experiment. For cell activation, the cells were concentrated in 50 μL imaging buffer and added to OKT3-coated coverslips (in a 35 mm dish, 5 μg/mL) for 10 min at RT. Afterwards, 2 mL of 4% PFA were added carefully and fixed the cells at 4C for 20 min and RT for another 20 min. After washing with PBS buffer 3 times, fixed cells were blocked with 10% goat serum in IF buffer (0.02% Triton-X 100, 0.005% Tween 20, 0.01% Bovine Serum Albumin prepared in PBS) at RT for 1 h. Primary antibody against pERK (Invitorgen, #MA5-15174) or primary rabbit monoclonal antibody against phosphoY171 of short LAT isoform (ab68139) was diluted 200 times in IF buffer with 10% goat serum and incubated with the cells at 4°C overnight. After washing 3 times with IF buffer, AlexaFluor647-labeled secondary antibody (goat anti-rabbit, #A27040, 2 μg/ml) in IF buffer was incubated with the cells at RT for 1 h. After washing 3 times with IF buffer and twice with PBS, the cells were imaged under epifluorescence microscope and TIRF microscope as mentioned above. For pERK induction by crosslinking Thy1, Jcam2.5 cells were mixed with Grb2-scarlet expressing Jurkats. After labeling with Thy1 primary antibody, cells were plated on secondary antibody-coated coverslips for 20 min before fixation and imaging.

*GPMV isolation and measurement of raft partitioning coefficient ($K_p$) and Tmisc:* GPMVs were produced, and partitioning quantified essentially as previously described (Sezgin, Kaiser et al. 2012, Levental and Levental 2015). Briefly, RBL or Hela cells were transfected with protein constructs (raft-TMD, nonraft-TMD, CD45-TMD, and GPI-GFP) or stained with fluorescent probes (chol-PEG-FITC, chol-PB+avidin, DOPE-PEG-KK114, DSPE-PEG-KK114), or primary and secondary antibody crosslinked Thy1 proteins as above. Cells were then counter-stained with disordered (nonraft) phase markers (either 10 µg/mL F-DiO, F-DiI or DiD, depending on fluorophore) for 5 min at 4ºC. Then GPMV production was induced by incubating cells in GPMV buffer (100 mM NaCl, 2mM $CaCl_2$, 2 mM dithiothreitol, 25 mM paraformaldehyde, pH 7.4) at 37°C for 1 h. Phase-separated GPMVs were imaged by epifluorescence microscope after cooling down to ~10°C using Peltier stage (Warner Instruments). K$p$ was measured based on the formula of $Kp = (I_{Lo} - I_0)/ (I_{Ld} - I_0)$, with Ld markers defining the nonraft (marker rich) and raft (marker poor) phases. $I_{Lo}$ and $I_{Ld}$ are the intensities in the probe channel from raft and nonraft regions. $I_0$ is background intensity. $T_{misc}$ was measured using the same method as for GUVs described above.

*Quantification of relative enrichment of GPI-anchors (GPI-GFP, Thy1-1°Ab, Thy1-2°Ab), raft-TMD, and nonraft-TMD under Grb2 condensates.* Image analysis was performed using ImageJ. Cells co-expressing Grb2 and either raft-TMD, nonraft-TMD, or GPI anchors were captured as time series from the initiation of activation upon cells touching down on the coverslips. Grb2 condensates were observed very quickly after cells' interaction with substrate; however, these initial condensates were irregularly shaped and dynamic, rapidly changing shape and size as the cells activated and spread. However, within 5 min, the macroscopic dynamics of Grb2 condensates slowed and came to a steady-state, characterized by similarly sized, approximately round condensates distributed throughout the bottom surface of the cell. All subsequent analyses were done on cells in this state. For quantification of membrane component enrichment under Grb2 condensates, Grb2 signal was thresholded and regions of interests (ROIs) were created. The mean intensity of the membrane probes within the ROIs were then compared to at least 5 ROIs with regions that contained the membrane probes but never had a Grb2 condensate throughout the image series. Shown in enrichment graphs (Fig 2F and Fig 3D) are the ratios of the mean intensity of the membrane probe under a Grb2 condensate versus non-condensate regions.

*Quantification of dynamics of Thy1-2°Ab clusters in Jurkat on OKT3/ICAM1 coated surface.* Time series (steps = 0.7-1.5 sec) of coverslip-plated Jurkat cells with 2°Ab-crosslinked Thy1 were acquired and analyzed with the TrackMate plugin of ImageJ (ref: PMID 27713081). The Difference of Gaussian (DoG) detection algorithm was used with a blob diameter set to 250 nm and an initial threshold of 5. In the tracking portion, the maximum allowable distance between frames was 0.3 μm with no gap closing across time steps. Only tracks containing at least 10 consecutive localizations were analyzed to avoid flittering noise. Consecutive time steps with <0.05 μm/sec velocities were interpreted as stalled and the percentage of any individual track spent stalled was quantified. Tracks with < 20% stalled were termed 'moving'; 21-79% stalled were 'start+stop'; >80% stalled were 'stalled'.

*Quantification of pERK intensity as a function of LAT intensity on PM*: pERK intensity was quantified as the mean intensity of pERK in each cell. LAT intensity on PM was determined from the peak value after drawing a line scan across each cell, similar to the method of GPMV *Kp* quantification described above. Both pERK and LAT intensity were subtracted by the background intensity before plotting. Each single point in the figure represents a single cell. Data representative of at least two independent experiments.

*Myriocin + Zaragozic Acid treatment of cells and quantification of condensate density:* Jurkat cells stably expressing Grb2-mScarlet were treated with 25μM myriocin and 5μM Zaragozic acid or DMSO (0.03 v/v%) for 2 days. 18 hr prior to imaging, FBS was removed from the medium (and drugs maintained in the medium). Cells were plated on OKT3-coated coverslips and activated at 37°C for 10 minutes. Then the cells were fixed with 4% paraformaldehyde for 20 minutes at 4°C then 20 minutes at RT and imaged via TIRF microscopy. Grb2 condensate density was determined by counting the number of condensates per 16 $\mu m^2$ areas of the cell.


Honigmann, A., V. Mueller, S. W. Hell and C. Eggeling (2013). "STED microscopy detects and quantifies liquid phase separation in lipid membranes using a new far-red emitting fluorescent phosphoglycerolipid analogue." Faraday Discuss **161**: 77-89; discussion 113-150.

Levental, I., D. Lingwood, M. Grzybek, U. Coskun and K. Simons (2010). "Palmitoylation regulates raft affinity for the majority of integral raft proteins." Proc Natl Acad Sci U S A **107**(51): 22050-22054.

Levental, K. R. and I. Levental (2015). "Isolation of giant plasma membrane vesicles for evaluation of plasma membrane structure and protein partitioning." Methods Mol Biol **1232**: 65-77.

Levental, K. R., J. H. Lorent, X. Lin, A. D. Skinkle, M. A. Surma, E. A. Stockenbojer, A. A. Gorfe and I. Levental (2016). "Polyunsaturated lipids regulate membrane domain stability by tuning membrane order." Biophys J **110(8)**: 1800-1810.

Lorent, J. H., B. Diaz-Rohrer, X. Lin, K. Spring, A. A. Gorfe, K. R. Levental and I. Levental (2017). "Structural determinants and functional consequences of protein affinity for membrane rafts." Nat Commun **8**(1): 1219.

Sezgin, E., H. J. Kaiser, T. Baumgart, P. Schwille, K. Simons and I. Levental (2012). "Elucidating membrane structure and protein behavior using giant plasma membrane vesicles." Nat Protoc **7**(6): 1042-1051.

Su, X., J. A. Ditlev, E. Hui, W. Xing, S. Banjade, J. Okrut, D. S. King, J. Taunton, M. K. Rosen and R. D. Vale (2016). "Phase separation of signaling molecules promotes T cell receptor signal transduction." Science **352**(6285): 595-599.

Zeno, W. F., K. E. Johnson, D. Y. Sasaki, S. H. Risbud and M. L. Longo (2016). "Dynamics of Crowding-Induced Mixing in Phase Separated Lipid Bilayers." J Phys Chem B **120**(43): 11180-11190.
